# Robust and Reliable *de novo* Protein Design: A Flow-Matching-Based Protein Generative Model Achieves Remarkably High Success Rates

**DOI:** 10.1101/2025.04.29.651154

**Authors:** Junyu Yan, Zibo Cui, Wenqing Yan, Yuhang Chen, Mengchen Pu, Shuai Li, Sheng Ye

## Abstract

Generative models have achieved significant progress in the field of protein design, particularly in the generation of tertiary structures. However, they still face several challenges, such as balancing the designability and diversity of the generated structures, generating specified structures under highly constrained conditions, and the most challenging aspect, direct functional design such as motifs and binders. We present OriginFlow, an efficient protein generative model that combines Stochastic Differential Equation (SDE) models and flow models based on flow matching. The generated structures exhibit state-of-the-art (SOTA) levels of designability, diversity, and novelty. Moreover, this model demonstrates outstanding performance in functional design aspects such as motifs, binders, and symmetric motif scaffolding. We performed binder design for multiple targets, and all results showed excellent affinity performance in AF3 complex predictions and MD experiments. Wet-lab validation was conducted on PD-L1, RBD, and VEGF targets, achieving 90% expression, solubility, and affinity. These results are significantly superior to all current AI binder design algorithms.

## 1 Introduction

Proteins are crucial molecules that execute most of the biological functions necessary for life, making their design essential for applications in biotechnology and medicine. However, designing proteins is an extremely complex task that traditionally relies on natural evolution, a process that spans billions of years.[1–3] The field of computational protein design aims to accelerate this process by using automated methods to design functional proteins within a programmable framework. This approach can significantly speed up the development of novel proteins for scientific research and practical applications.[4, 5]

Recent decades have witnessed remarkable advancements in protein structure prediction, largely driven by the application of deep learning methods. Notably, deep learning models such as AlphaFold2[4] and RoseTTAFold[6] have achieved unprecedented accuracy in predicting protein structures by utilizing translational and rotational equivariant networks for end-to-end modeling of sequential and three-dimensional data. These breakthroughs have laid a strong foundation for further developments in the field.

Building on these advancements, the focus has expanded to protein structure generation. Generative models, particularly denoising diffusion probabilistic models (DDPMs)[7] and score-based stochastic differential equations (Score SDEs)[8], have shown great promise in this area. These models have been extensively applied to de novo protein structure generation, achieving significant progress. The ability of these models to generate diverse and novel protein structures has opened new avenues for designing proteins with specific functional properties.[1, 9–11]

Despite recent advances, low success rates—especially in wet-lab validation—remain a major bottleneck in computational protein design, particularly for *de novo* binder design. Most existing models struggle to generalize beyond simple monomer structures, with performance dropping as residue length increases. They also often lack the ability to perform conditional generation under functional constraints (e.g., motif or binder design), and tend to over-optimize for structural stability at the cost of functional flexibility.

Fundamentally, protein structure generation is a challenging 3D task, requiring efficient modeling of atomic coordinates within a vast sampling space. Compared to discrete 2D domains like images, protein design demands significantly more computational memory and is highly sensitive to structural errors. Additionally, the variation in sequence lengths necessitates models with strong extrapolation capabilities, while the framework must remain extensible across diverse design tasks.

To address these challenges, we introduce **OriginFlow**, a generative model that integrates Flow Matching with unified SDE and ODE training, drastically reducing training complexity. OriginFlow employs a flexible transformation framework between SE(3) and ℝ ^3^, enabling efficient sampling and robust structure representation. Its architecture supports both global and local feature updates, achieving strong extrapolation and scalability to long sequences. OriginFlow achieves state-of-the-art results across monomer, complex, and functional conditional design tasks, with performance validated by both molecular dynamics and wet-lab experiments.

**Most importantly**, OriginFlow achieves an unprecedented success rate of over 95% in binder protein design, significantly outperforming existing protein generative models. In contrast, recent tools like AlphaProteo report wet-lab success rates ranging from 5% to 80% across different targets (e.g., 33% for VEGF). Our method consistently demonstrates 89% success in expression, solubility, and affinity in *in vitro* validation across three different targets. Specifically, for the VEGF target, 12 binders were successfully expressed and soluble, with the exception of one binder (VE2), which formed precipitation after overnight incubation. The remaining 11 binders exhibited affinities in the 10^*−*6^ M range. Additionally, 8 binders targeting **CovidRBD** and **PDL1** were all successfully expressed, soluble, and showed affinities of 10^*−*6^ M in SPR assays—without any biological post-processing. This underscores **OriginFlow**’s exceptional ability to generate realistic, high-affinity binders with unmatched reliability.

If high binder success rates can be consistently achieved *in silico*, this would dramatically reduce reliance on costly high-throughput screening. Instead, low-throughput validation could suffice, accelerating discovery pipelines, cutting experimental costs, and significantly increasing the efficiency of drug development. The impact on both academic research and pharmaceutical innovation could be transformative.

## 2. Results

### 2.1. An overview of OriginFlow

OriginFlow is a generative model based on the Optimal Transport Flow Matching (OTFM) framework(Methods). The model represents protein backbone structures as frames consisting of rotation matrices in SO(3) space and translation vectors in ℝ^3^ space like previous studies[10, 12, 13]. Fig. 1 A During training, the model learns to reverse the optimal transport path from the prior space *x*_0_ to the data space *x*_1_ (a linear path in ℝ^3^ space and a geodesic path in SO(3) space). The model calculates the loss as the difference between the reverse structure and the native structure in both ℝ^3^ and SO(3) spaces, while also constructing a complete mapping of the frame structure in ℝ^3^ space using the ℝ^3^ covariance matrix *z*.(SI C) This *z* serves as an approximate latent variable to calculate the Evidence Lower Bound (ELBO), ensuring the model’s generalization capability. These loss functions drive the model to converge on designable protein backbone structures along the fastest vector field direction at each time step.(SI B.2)

**Fig. 1:**
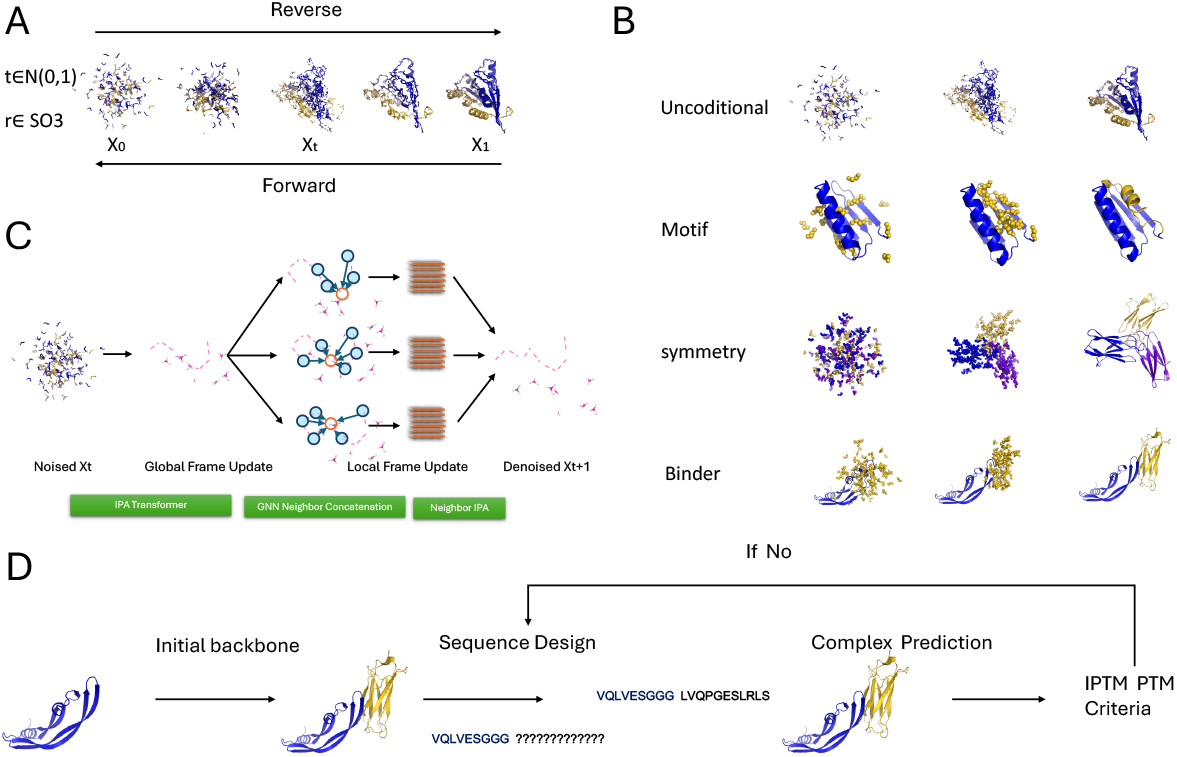
Structure generation using OriginFlow. **A**. Overall overview of OriginFlow. Protein structures are represented as combined vectors in both ℝ ^3^ space and SE(3) space. The model samples initial random structures by drawing from a Gaussian distribution in ℝ ^3^ and uniform distribution on the sphere in SO(3). The denoising network learns to reverse the noise process, enabling structure generation through reverse dynamics. **B**. OriginFlow supports a variety of generative tasks, including unconditional structure generation, motif scaffolding, symmetric assembly generation, and binder design. **C**. Schematic of the network architecture. The noisy initial structures are first passed through a global Invariant Point Attention (IPA) module for initial backbone generation. Then, local frames are constructed from structural neighbors to aggregate features, followed by another IPA module to update the local frame structures. **D**. Binder generation protocol. A fine-tuned model is used to generate the initial backbone structure. The target region is fixed while the binder region is sequentially filled. Structure prediction is performed using ESMFold or AlphaFold3. If the predicted interfacial pTM and pTM scores are both above 0.8, the generation is considered successful. Otherwise, the sequence is further refined until the criterion is met.

During sampling, the solution of the FM framework can be expressed as a constant velocity ODE[14], allowing each residue to iteratively follow the vector field direction in both ℝ^3^ and SO(3) spaces. Through continuous small iterations, the model gradually reaches the sample space to generate designable structures. The FM framework can also be extended to SDE and Probability Flow ODE(Methods)., enabling sampling in the pure ℝ^3^ covariance matrix space using their solution forms, thus efficiently incorporating common SDE/ODE sampling methods without dealing with the complex forms of SDE in SO(3) space.

The overall model comprises an initial global structure update module using invariant-point-attention[4] and a local detail update module GA (GNN and Local Attention) combined.Fig. 1 C The transformer part uses Relative Position Encodings (ROPE) [15] to enhance extrapolation capability. After obtaining the initial structure, the K-nearest neighbors (KNN) topology is determined based on the distances between Ca atoms, aggregating the K-nearest information around each point, and updating the central point’s structural information using a lightweight version of IPA.(SI D.2)

The data is sourced from the RSCB[16], filtered using sequence identity clustering at 30%. We first train a base model capable of generating good monomers and complexes, randomly running 50% of the time during training to obtain initial predicted structures for self-conditioning(SI E). Many studies have emphasized the importance of incorporating symmetries of the target system into the flow model. Therefore, we normalize the overall protein structure using the zero center of mass (CoM) subspace of ℝ^3^, which also helps reduce distance discrepancies caused by rotations. Additionally, pre-alignment is used to accelerate the training process[17, 18]. For the binder task, we fine-tune using protein-protein complexes from the PDBbind database.(SI E.2)

After fine-tuning, the model can be directly applied to various application scenarios, such as unconditional multi-chain generation, motif scaffolding, symmetric structure generation, and binder design Fig. 1 B. In the binder design phase, in addition to the model, we have also developed a set of protocols, as shown in Fig. 1 D. A detailed description is provided in SI J and Binder Design.

### 2.2. Unconditional protein monomer generation

As shown in Fig. 2 A,B, OriginFlow can generate proteins of any length, and even when the generated sequence length far exceeds the training length (384 aa), the model still performs well. The generated protein structures have a relatively high proportion of secondary structure (80%), higher than RFdiffusion and Chroma.SFig. 5 C As the length increases, the model improves stability by reducing the proportion of coil and strand regions and increasing the proportion of helix regions. SFig. 5 B,C. We used ESMFold[19] to fold the sequences filled with these generated structures (using ProteinMPNN[20]) for refolding analysis. As shown in Fig. 2 B, within 500 aa, both OriginFlow and RF have a mean pLDDT above 80, and most TM-scores are above 0.9, indicating that the generated structures can be easily filled with sequences and folded with high similarity. Additionally, both OriginFlow and RFdiffusion can control RMSD within 5 Å, and even within 2 Åfor sequences shorter than 400 aa. In the length range of 40-600 aa, evaluated by the medians of pLDDT, RMSD, TM, and pAE, OriginFlow (pLDDT=80.8, scTM=0.9, scRMSD=1.9) approaches RF (pLDDT=86.8, scTM=0.97, scRMSD=1.0) in high designability, far surpassing Chroma (pLDDT=57.35, scTM=0.65, scRMSD=6.9). We also calculated the decay and correlation of various metrics with length for the three models, as shown in SFig. 3. OriginFlow (*R*_*pLDDT*_ = *−*0.52, *R*_*T M*_ = *−*0.35, *R*_*rmsd*_ = 0.52) has lower correlation with length compared to RF (*R*_*pLDDT*_ = *−*0.61, *R*_*T M*_ = *−*0.47, *R*_*rmsd*_ = 0.54),SFig. 4 indicating weaker decay with length and better extensibility.

**Fig. 2:**
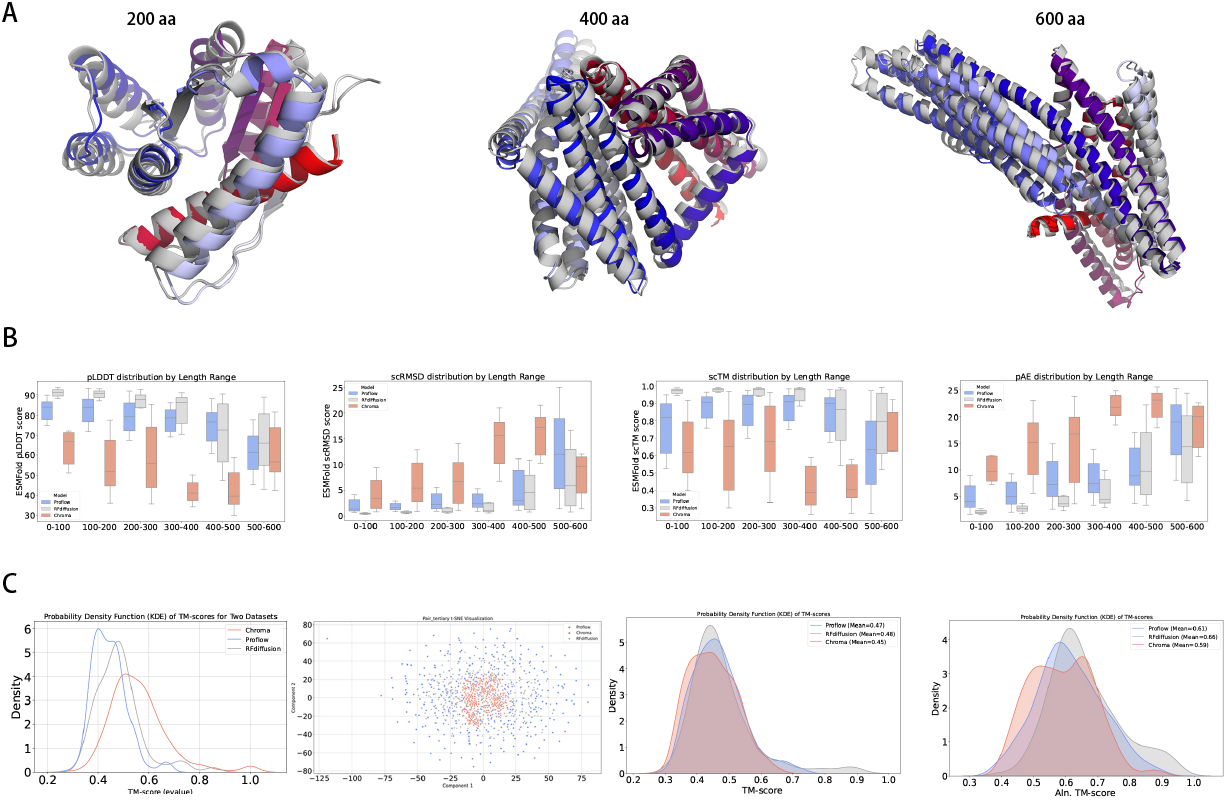
Performance of OriginFlow in Unconditional Generation. **A**. OriginFlow can generate proteins of various lengths, and it performs well even when generating lengths far exceeding the training lengths. The gray structures are generated structures, while the colored structures are ESMfold predicted structures. The full-sequence alignment r.m.s.d (Å) of the three structures are 1.2, 1.7, and 2.3, respectively. **B**. Refolding experiments were conducted on OriginFlow, RFdiffusion, and Chroma models. The plots from left to right show pLDDT, r.m.s.d, tm-score, and pAE versus length. It can be observed that OriginFlow is slightly inferior to RFdiffusion but significantly better than Chroma across different length ranges. Additionally, the decline with increasing length is relatively moderate for OriginFlow. **C**. Pairwise TM-score comparisons were made for structures generated by each model, and their probability density plots were drawn. OriginFlow is overall on the far left, indicating the lowest overall similarity and the highest diversity of generated structures. The mean pair TM-scores for OriginFlow, RFdiffusion, and Chroma are 0.45, 0.48, and 0.56, respectively. **D**. Distribution of TM-scores between generated structures from the three models and CathS40 structures. TM-score represents the overall average comparison result, and Ala. TM-score is the average TM-score for the aligned residue parts. Structures generated by Chroma have the lowest similarity to known models under both evaluation scores, followed by OriginFlow, with RFdiffusion showing the highest similarity (highest TM-score) to known structures. **E**. TM-scores categorized by different sequence lengths. As the length increases, the similarity decreases for all models, but RFdiffusion shows abnormally high similarity in short sequences.

The structures generated by OriginFlow are highly diverse. We calculated the pair-wise similarity (TMscore) within the generated structure sets of these three models. As shown in Fig. 2 C, OriginFlow has the lowest mean (0.45 vs RF 0.48 vs Chroma 0.52). We further mapped this pairwise similarity using T-SNE to show their spatial cluster distance structure, as shown in Fig. 2 D. The structures generated by OriginFlow have greater distances between each other in the T-SNE mapping space [±50], while Chroma has only [±20].

Furthermore, to evaluate the novelty of structures generated by each model, we used Foldseek to find the most similar structure with the highest hit score in CATHS40 for each design (taking the maximum TMscore), as shown in Fig. 2 E. OriginFlow’s TM-score (overall) and alaTM-score (aligned residue TM-score) were between Chroma and RF diffusion (OriginFlow 0.61, Chroma 0.59, RF diffusion 0.66). We analyzed the novelty by length, and it can be seen that as the length increases, the similarity gradually decreasesSFig. 5A, Right. However, RF shows high similarity to existing structures in short sequences, while OriginFlow and Chroma exhibit higher novelty.

Overall, our method consistently generates highly designable structures across long sequence ranges, with a good secondary structure distribution and substantial diversity and novelty. Compared to the current SOTA, RF diffusion performs better in designability but lacks innovation, while Chroma exhibits the highest structural novelty but suffers from uneven sampling distribution and lower designability.

### 2.3. Functional-motif scaffolding

The purpose of structural design is to serve functional design, so we further investigate the performance of the model in scaffolding protein structural motifs. We fine-tuned the model so that during training and inference, the backbone atoms of the motif regions are preserved, while other regions are randomly sampled from the sampling space. During the Motifs-generating iterative stages, only the non-motif parts are updated. We used a set of 17 novel protein structures as a benchmark test[10, 21]. These proteins, containing motifs with various functional sites such as viral epitopes, receptor traps, ligand interfaces, small molecule binding sites, and enzyme active sites, were released after the data collection cutoff date for the training set, ensuring no overlap. For each motif, we generated 100 structures and used ESMFOLD[22] to predict the structures for the completed sequences. If the overall pLDDT is greater than 70 and the motif RMSD (i.e., the RMSD of atomic positions between the motif residues in the predicted structure and those in the pre-defined motif structure) is less than 1.0Å, the motif design problem is considered solved.

The results show that our model solved 16 out of the 17 problems, which is the highest among RF diffusion [10], EVODIFF [23], and PVQD [21]. (Fig. 3A) Despite varying success rates for different motif designs, OriginFlow consistently achieves an RMSD error below 2Å in 90% of cases (Fig. 3B). In scenarios where motifs are located in multi-chain regions (Fig. 3D) or in multiple motif regions (Fig. 3E, F, G), OriginFlow always manages to design high-quality scaffolding. 3ixt, 2kl8, and 1bcf are representatives of single-chain multiple-motif cases, with individual motif regions typically ranging from 5 to 20 residues in length and inter-motif distances generally between 15 to 30 residues. OriginFlow demonstrates high-confidence scaffolding in these cases. The motif region of 6vw1 consists of three motifs from two chains, and we successfully designed it as well, achieving an RMSD of 0.74Å and a pLDDT of 86.61 (Fig. 3D).

**Fig. 3:**
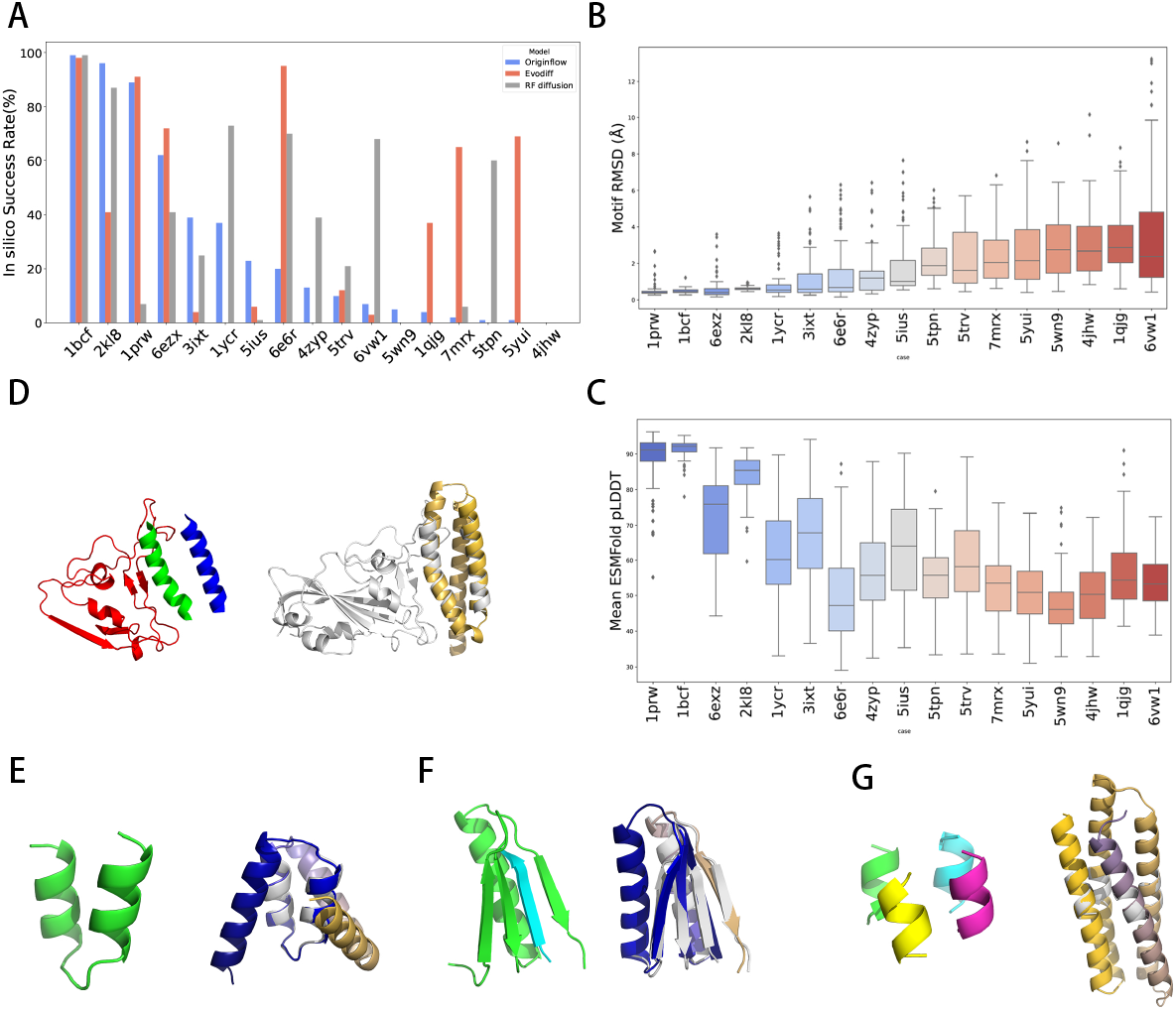
Motif-Scaffolding using OriginFlow. **A**. OriginFlow demonstrates higher success rates in motif design on a test set of 17 benchmarks. A design is defined as successful if the motif r.m.s.d is less than 1Å, and the overall ESMfold pLDDT is greater than 70. **B**. The distribution of rmsd values for different motif design cases. **C**. The distribution of ESMfold pLDDT values for different motif design cases. **D**. The 6vw1 double-chain motif design case. The left image shows the fixed motif, with red on the E chain and blue and green on the A chain. The right image shows the AF3 prediction of the design case, with the motif part in gray and the scaffolding part in color. Metrics: RMSD/pLDDT 0.74/86.61. **E. F. G**. Single-chain single motif 3ixt_88, single-chain double motifs 2kl8_4, and single-chain triple motifs 1bcf_88. OriginFlow performs well with different numbers of motifs. The left images show the motif regions, and the right images show the designed motif-scaffolding in color. Metrics: RMSD/pLDDT 0.47/93.12, 0.66/92.13, 0.56/90.94.

Additionally, following the RF diffusion approach, we discretized the scaffolding length of some motifs (generating short, medium, and long scaffolds), resulting in a total of 25 test experiments, of which we solved 24. (SI Fig. 6) Although RF diffusion shows a higher average success rate, it tends to fail more easily in some long-tail cases. This could be related to the lower diversity in RF diffusion model designs.

### 2.4. Geometry-Constrained Generation

In de novo generation tasks, it is sometimes necessary to impose prior structural constraints on specific regions or modify the structure of certain areas [9]. The primary structural constraint is the secondary structure of proteins. We fine-tuned the model to generate structures based on specific secondary structure inputs. Referring to representative PDB secondary structure sequences from the three CATH classifications—main alpha, main beta, and alpha-beta—we designed structures for 27 CATH architecture types. SI:Tab2. We designed 20 structures for each architecture and compared the native secondary structure sequences with the generated secondary structure sequences. We used precision, accuracy, recall, and F1 score to measure the consistency between them Fig. 4A. The overall consistency (F1 Score) ranged between 80% and 95%, performing well across sequences ranging from 100 residues (1xmkA00) to 700 residues (5kisB00) Fig. 4C. The highest performance was observed for Main Alpha, averaging 93%, followed by Main Beta at around 81%, and AlphaBeta at approximately 85%. The primary factor contributing to this difference is the amount of Coil in the native sequence; fewer Coil regions result in higher accuracy Fig. 4B, which is reasonable as Coils primarily provide flexibility.

**Fig. 4:**
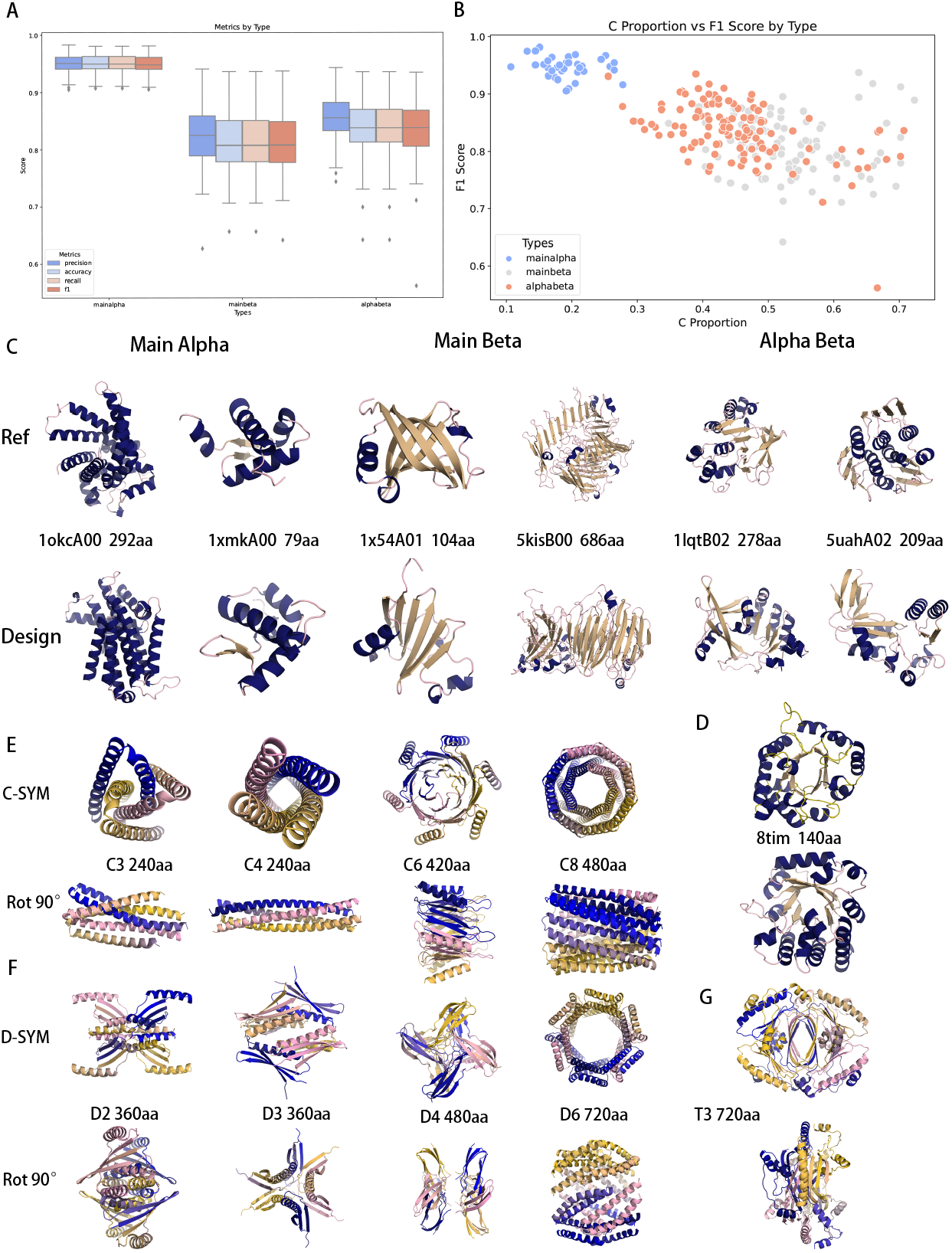
Geometry-Constrained Generation. **A**. Conditional generation accuracy under different main types of secondary structures. **B**. Relationship between secondary structure consistency and the proportion of Coil in the input sequence. The lower the proportion of Coil, the higher the accuracy. **C**. Using the secondary structure sequence of the protein in the reference row as input, the generated structure in the design row matches the input type regardless of the sequence length. **D**. We generated a protein structure based on the secondary structure sequence of 8tim, successfully obtaining the TIM structure. **E**. Cyclic symmetric polymers, with total lengths ranging from 240aa to 480aa.

Additionally, we performed reference designs on some classical structures, such as the TIM-barrel structure (8tim). The designed structures showed 88% to 92% consistency with the original secondary structures and could all form TIM-barrel structures Fig. 4D. This demonstrates that our secondary structure constraints are highly accurate and that the model is sensitive to secondary structure inputs.

Another important structural constraint is the design of symmetric polymers. Symmetric structures are very common in many functional complexes, especially in drug pockets, vaccines, or nanocages. However, due to the longer sequences of symmetric structures, relatively less data, and high precision requirements, it is difficult to accomplish this through model fine-tuning. Therefore, we adopted a method of guiding the gradient directly during the sampling process to achieve symmetric structure design. This method references RFdiffusion and Chroma, where after obtaining an *X*_0_ state at each reverse step, we select a portion of the subunit for rotational symmetry replication to obtain 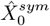, and then calculate the iterative gradient. In this way, during the multiple sampling processes, the structural vector evolves towards a combination of unconditional constraints and symmetric constraints, eventually achieving a symmetrically constrained structure.

We generated several cyclic symmetric samples, including C3, C4, C6, and C8, with single-chain lengths ranging from 60 to 120 Fig. 4E. For dihedral symmetric bodies, we designed d2, d3, d4, and d6 Fig. 4F. Additionally, we designed a T3 symmetric polymer with a total sequence length of 720 Fig. 4G.

### 2.5. Binder Design

Binder design is one of the most critical topics in de novo protein design and represents a grand challenge in the field of protein design. Traditionally, this has been approached using physical methods, such as the continuous mutation and computational iteration approach of Rosetta. Recently, generative models have provided new possibilities for directly generating high-affinity binders and have shown promising results in some models.

We performed *de novo* binder design targeting several therapeutically relevant proteins, including PD-L1, MDM2, SARS-CoV-2 Spike RBD (SRC2RBD), GLP-1R, GP120, KRASG12, and VEGF. Among these, we selected PD-L1, SRC2RBD, and VEGF for *in vitro* experimental validation.

#### 2.5.1. Design binder proteins against given targets

We regard binder generation as a conditional generation model, where *P* (target, binder) = *P* (target) *× P* (binder | target). Based on the foundational model, we fine-tuned it using complex data and PDBBind data. We first extracted interface residues from complexes (not exceeding 384 residues), then randomly masked one end and initialized it as noise. The remaining structure and sequence served as the target chain reference, and their features were computed and fed into the model. During the reverse iteration process, the model gradually restored the disturbed noise into the binder structure. After obtaining the complete binder structure, we filled the binder sequence based on the overall structure and target chain sequence.

In our approach to target-specific binder design, we developed a protocol to refine the binder design process. We first employed a foundational model based on target structure and sequence to perform a reverse iteration process, generating the initial scaffold. Subsequently, we input the generated binder scaffold and target into MPNN using binder mode to populate the binder sequence. We then conducted preliminary screening using ESMFold. To ensure structural diversity, we selected the most structurally diverse candidates from the top 5-10 cases and submitted them to AlphaFold3[5] (AF3), which has demonstrated robust capabilities in complex prediction.

When designs achieved both iPTM and PTM values exceeding 0.8, we considered the design complete. Otherwise, we continued the process using the AF3-predicted binder structure as a foundation, repeating the sequence population and prediction validation steps. According to our experience, no more than two AF3 iterations were required to obtain the final results.

We initially applied this protocol to design binders for PD-L1, GP120, GLP-1R, G12D, and SC2RBD, with specific details provided in the Supplementary Information. SI:Tab3, SI:Tab4

#### 2.5.2. Physicochemical characteristics of designed binders

A total of 17 selected designs were recombinantly expressed in *E. coli*. SDS-PAGE detection showed that all designed proteins were in the soluble fraction of the lysate (SI SDS). A subsequent purification via nickel affinity chromatography resulted in rough yields ranging from 0.8 to 3.2 mg per liter LB culture. OriginFlow’s designs showed high solubility and high success rate in recombinant expression. This performance surpasses that of proteins designed by other widely used AI methods. Circular dichroism (CD) spectroscopy was further applied to characterize the folding quality of the designs, as well as their thermal stability. According to the CD spectrum collected, all designed binders had folded correctly, showing strong signals of proper secondary structures as in their designed structures Fig. 5G. The melting temperatures (Tm) of 4 designs varied from 60 to 85 °C, exhibiting high thermal stabilities in solution. The other 7 designs showed no obvious changes in CD absorbance at 222 nm during the temperature range of 20 to 90 °C, suggesting that their Tm could be higher than 90 °C. The high thermal stability of OriginFlow’s designs is consistent with the findings unveiled in previous AI protein design studies.

**Fig. 5:**
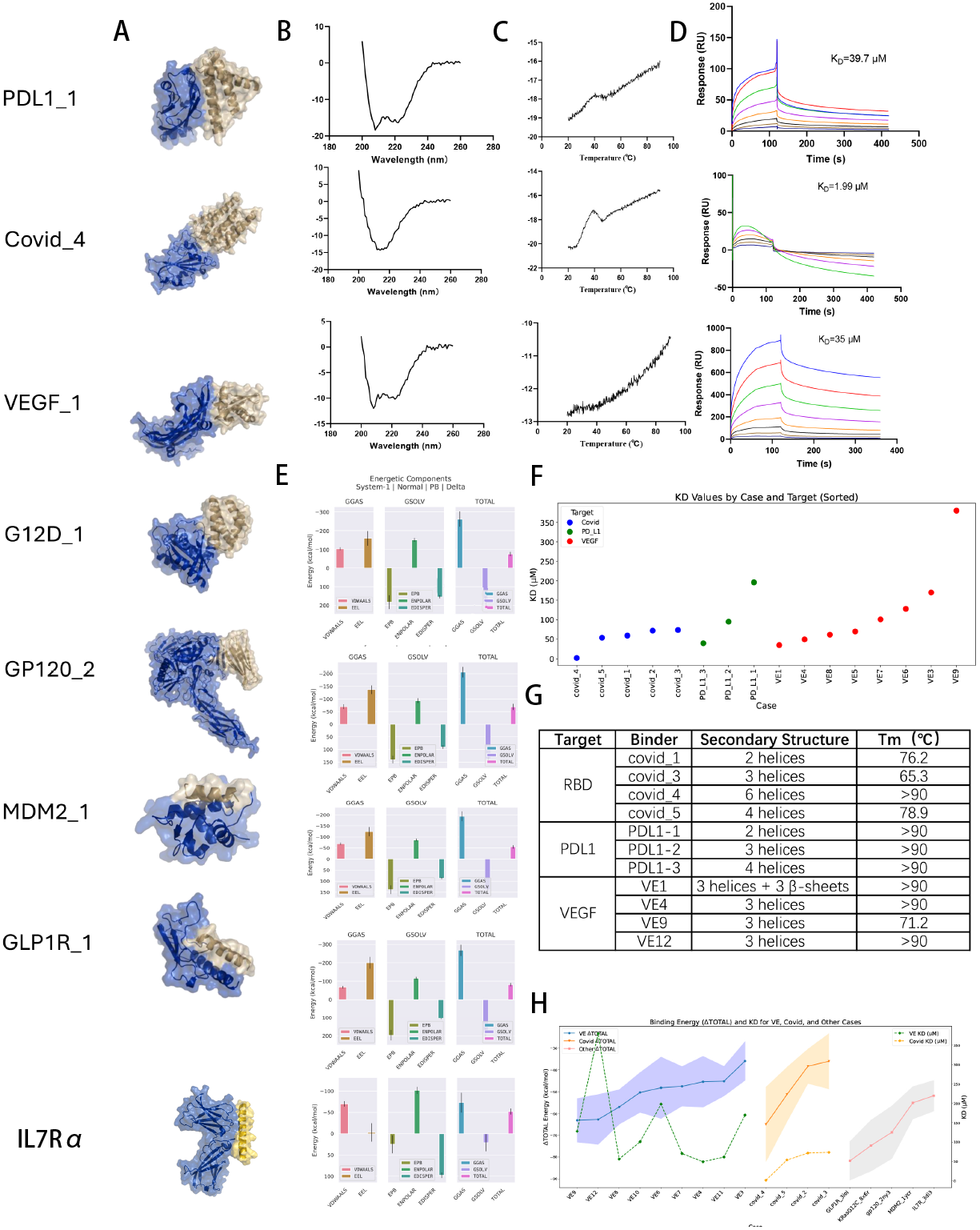
Binder design results of OriginFlow. **A**. Schematic representation of binder designs targeting different epitopes, where the blue indicates the target and the yellow represents the binder. Each target shows one design result. **B**. Circular dichroism (CD) spectra from wet experiments, demonstrating that the experimental CD results are consistent with the secondary structure predictions. **C**. Thermal melt (TM) plots of different binders. **D**. Surface plasmon resonance (SPR) results of different binders, with KD values in micromolar (*μM*) units. **E**. Binding energies calculated using GROMACS for several targets, showing negative binding energies (ranging from −50 to −100 kcal/mol) for five binders, indicating strong affinity. **F**. Experimental SPR binding affinity results for three categories of targets, sorted by binding strength. Except for VE2, which showed aggregation, the other binders had KD values between 10^*−*6^ and 10^*−*4^ M, qualitatively confirming their affinity. **G**. Thermal stability and secondary structure composition of different binders, measured by circular dichroism at 222 nm. The CD signal at 222 nm primarily reflects *α*-helical content. All binders exhibited good thermal stability, as evidenced by their melting temperatures (*T*_*m*_). Several binders showed *T*_*m*_ values exceeding 90°C, which is the upper detection limit of the instrument. **H**. MD-calculated binding energies for several wet-lab-tested binders, plotted on the left axis, and their corresponding KD values, plotted on the right axis. It is observed that VEGF binders have relatively high MD values, while the COVID-related binders are slightly lower. KD values also show that COVID-related binders generally exhibit lower affinity compared to VEGF binders. The MD values for the remaining targets are lower than both VEGF and COVID categories, suggesting that although no wet-lab experiments were performed for these, their binding affinities are likely to be lower, likely reaching KD levels around 10^*−*6^ M.

#### 2.5.3. Extremely high success rate of binder designing

Due to the limited number of samples to be tested, we bypassed the conventional high-throughput or medium-throughput evaluation approaches and instead conducted individual affinity measurements on each sample directly. To quantitatively assess the binding affinities between OriginFlow’s designs and target proteins, Surface Plasmon Resonance (SPR) assays were performed to accurately measure the dissociation constant *K*_*D*_ of each binder-target pair. For the 20 designs, the *K*_*D*_ values ranged from 2.0 to 200 *μ*M. Among them, 7R-34 and its target SARS-CoV-2 RBD showed a *K*_*D*_ value of 2.0 *μ*M. It is noteworthy that all 17 designs exhibited binding affinities to their designed targets, leading to a 90% success rate. To the best of our knowledge, OriginFlow’s extremely high success rate in binder designing makes it remarkably unique. Although the moderate binding affinities may limit any direct uses of the designs, the robust success rate dramatically reduces the time and cost in the “binder-finding” stage and provides solid foundations for further affinity maturation or structure-based optimization.

## 3. Methods

Topical subheadings are allowed. Authors must ensure that their Methods section includes adequate experimental and characterization data necessary for others in the field to reproduce their work. Authors are encouraged to include RIIDs where appropriate.

### Optimal transport flow matching model

(SI B.1)Flow Matching does not require a step-by-step simulation process for training, as DDPMs and SDEs do. Instead, it directly regresses vector fields. On one hand, Flow Matching focuses on the direct probabilistic distribution transfer between the perturbation space and the target space, which has relatively low transport costs [24], significantly reducing computational costs, especially in high-dimensional data processing. On the other hand, the objective of Flow Matching is to obtain a vector field *u*_*t*_(*x*) that can generate the target probability density path *p*_*t*_(*x*). Therefore, we can directly construct relatively simple and easy-to-train vector fields (VFs). Here, we use a linear interpolation path between the source distribution and the target distribution. This path differs from the VE path used by DDPM[7, 8] and the VP path used by SDE[8], which are deterministic transition probability density paths controlled by a single parameter [1, 14]. These optimizations significantly enhance training efficiency [12, 25, 26].

### Inverse Process

We have constructed two representations for protein structures: the covariance matrix and the frame. The frame process represents atoms with a translation and a rotation matrix, similar to our forward process. The covariance matrix is represented by the C-alpha atoms and the residue atomic coordinates centered on the C-alpha atoms. Under the frame, we can directly construct the CVODE reverse sampling process, iterating separately on the translation and rotation matrices. Sampling can also be performed under the covariance matrix using SDE and PFODE, and we have proven their mathematical equivalence.

### Model Architecture

Our model consists of six transformers with structural attention and three layers of GNN-transformer. The former primarily generates the global structure, while the latter uses the structural attention transformer for detailed updates on neighboring nodes. Rotational position encoding is added to the attention calculation, and length-based attention scaling is used to mitigate the entropy increase phenomenon of transformer length.

### Training

We use RSCB structural data with a resolution less than 2.6 and sequences shorter than 1000 in length, totaling 17k sequences. The structural data is segmented to a length of 384, mainly using the structural constraints composed of translation and rotation matrices, and adding a physical rationality representation loss (Violoss). We also constructed a latent ELBO constraint using the covariance matrix. For motif and binder tasks, we fine-tuned the data.

### Inference

We provide detailed descriptions of the inference settings for each task in the supplementary materials. The configurations are largely consistent across tasks, with the exception that the number of inference steps is set to 500 for all tasks except for the binder design task, which uses 1000 steps.

### Sequence Completion and Structure Prediction

We used the MPNN and ESMFOLD suite provided by Genie, following the same parameter configurations as in the original Genie setup. For AlphaFold3 predictions, we utilized the service provided by DeepMind.

### Purification of designed binder proteins and target proteins

All DNA sequences coding designs and targets were synthesized by Tsingke Biotechnology Co., Ltd. and subsequently cloned into the pET-28a vector. These constructs were then transformed into BL21 (DE3) (Tsingke, Beijing, China) and cultured in LB medium at 37°C with vigorous shaking. Protein expression was induced by adding 0.5 mM IPTG when the OD600 reached 0.8, followed by overnight culture at 16°C.

For purification of recombinantly expressed proteins, cells were harvested by centrifugation and resuspended in PBS. The resuspension solution was lysed by sonicating and then centrifuged at 12,000 *×* rpm for 30 min at 4°C. The supernatant was applied to NTA resin pre-equilibrated by PBS with 20 mM imidazole. The eluate was achieved with PBS containing 250 mM imidazole, which was then identified by SDS-PAGE and concentrated for further evaluation. The concentration of each protein sample was measured by the absorbance at 280 nm with a Nanodrop One (Allsheng Instruments, Hangzhou, China).

For purification of target proteins including SARS-CoV-2 RBD, programmed cell death ligand 1 (PD-L1) extracellular domain, and vascular endothelial growth factor (VEGF), the harvested cells were processed as described above. The pellet was resuspended in lysis buffer (20 mM Tris, 0.5 M NaCl, 0.1% Triton-100, 1 mM PMSF, pH 8.0), followed by sonication for 15 minutes and centrifugation to collect the precipitate. The precipitate was then washed three times with Buffer A (50 mM Tris, 0.1 M NaCl, 1 mM EDTA, pH 8.0) and once with distilled water.

The resulting pellet was dissolved in Buffer B (20 mM Tris, 0.5 M NaCl, 8 M Urea, pH 8.0) and loaded onto a pre-equilibrated Ni-NTA affinity column. Target proteins were eluted using a linear imidazole gradient (20–250 mM) in the same buffer. Eluted fractions were collected and analyzed by SDS-PAGE.

The purified target proteins were subjected to stepwise dialysis against Buffer B containing decreasing concentrations of urea (8 M, 4 M, and 2 M), with each step lasting at least 4 hours. A final dialysis was performed in urea-free Buffer B to complete protein refolding for subsequent experiments.

### Circular dichroism (CD) spectroscopy

Data were collected on a Chirascan Plus circular dichroism spectrometer (Applied Photophysics Ltd., North Carolina, United States). All samples were diluted to 0.2 mg/mL in PBS for measurements in the UV spectral range of 200–260 nm and for thermal denaturation analysis.

Thermal melt analyses were performed at 222 nm from 20 °C to 90 °C with a temperature ramp rate of 2 °C/min, and measurements were recorded at 0.2 °C intervals. All reported measurements were obtained within the linear dynamic range of the instrument.

### Surface Plasmon Resonance (SPR)

SPR data were acquired on Biacore T200 instruments (Cytiva) at room temperature. Target proteins were immobilized on individual channels of CM5 sensor chips by a standard amine-coupling approach. Briefly, the sensor surface was activated with a 7-minute injection of a mixture of 50 mM N-hydroxysuccinimide (NHS) and 200 mM 1-ethyl-3-(3-dimethylaminopropyl) carbodiimide (EDC). Target proteins at a concentration of 50 *μ*g/mL in 10 mM sodium acetate buffer (pH 4.0) were injected into individual channels on sensor chips at a flow rate of 10 *μ*L/min for 420 s. Then, the channels were blocked with 1 M ethanolamine (pH 8.0). Designed binder proteins were dissolved to 200 *μ*M in 1 *×* HBS-EP running buffer and then diluted 2-fold into 100 *μ*M, 50 *μ*M, 25 *μ*M, 12.5 *μ*M, 6.25 *μ*M, 3.125 *μ*M, and 1.56 *μ*M before injection. Data were analyzed with Biacore T200 Evaluation Software.

## 4. Discussion

We propose a novel application of FlowMatch for protein design, particularly for binder design. Simultaneously, we optimized the method from multiple aspects, including data, algorithm, and structural considerations, specifically tailored to protein-related tasks. Our approach demonstrates strong generative capabilities and favorable experimental characteristics. Notably, we have achieved an almost 90% success rate across all binder design cases tested so far. This indicates that our algorithm is capable of generating reasonable initial structures and effectively identifying near-optimal solutions during the iterative refinement process.

This represents a significant advancement in the field of *de novo* protein design. It suggests that we may be able to rapidly and cost-effectively generate large quantities of initial binder candidates, thereby circumventing the need for high-throughput screening traditionally required in binder development. If the success rate of *in silico* binder design can be maintained at a sufficiently high level, it opens up the possibility of validating results using only low-throughput experiments. This would allow for the identification of initial binders while drastically reducing experimental costs and increasing both the speed and success rate of the discovery process. Such improvements could have profound implications for the pharmaceutical research and development industry as a whole.

Moreover, there remains substantial room for optimization in the current algorithm. One major limitation lies in the separation between backbone and side-chain modeling, which necessitates the use of an additional MPNN module for sequence completion. We believe this is a key factor contributing to the binding affinities of our designed binders remaining in the micromolar range rather than reaching the nanomolar level. As shown in the supplementary materials, many residues in the final designed binders contribute negatively to binding affinity; these residues are retained primarily to preserve the original backbone conformation. If side-chain considerations could be integrated directly into the generative phase, we anticipate that binding affinity could be significantly improved.

## Supporting information

SICD

SISDS

SISPR

SITM

## Supplementary information

This manuscript includes 5 supplementary files that provide detailed experimental procedures and additional experimental results not included in the main text:

- **Supplementary Information (SI)**: Contains comprehensive descriptions of experimental details and supplementary experiments that support the findings presented in the main manuscript.
- **SI_SPR**: Provides detailed information on surface plasmon resonance (SPR) experiments, including binding affinity measurements, experimental parameters, and additional analyses.
- **SI_CD**: Contains circular dichroism spectroscopy data, methodology, and analyses that characterize the secondary structure of the studied molecules.
- **SI_TM**: Presents thermal melting experimental results, including temperature profiles, stability analyses, and methodological details.
- **SI_SDS**: Includes the complete set of electrophoretic gel images from SDS-PAGE experiments with detailed annotations and experimental conditions.

## Declarations

### Funding

This research is supported by the Beijing Municipal Science & Technology Commission grant [Z221100003522020] and the National Key Research and Development Program of China NO.2023YFF1203803, the National Natural Science Foundation of China No. 62272021.

## Conflict of interest/Competing interests

The authors declare that they have no conflict of interest or competing interests.

## Data availability

All training data used consists of publicly available datasets, and the methods used as well as the generated cases mentioned in the paper will be open-sourced at https://github.com/JoreyYan/Originflow.

## Materials availability

All materials used in this study are available from the corresponding author upon reasonable request.

## Code availability

All code used in this study will be made publicly available on GitHub upon publication.

## Author contribution

Junyu Yan developed the overall algorithm and wrote the manuscript. Zibo Cui conducted all wet lab experiments. Shuai Li and Sheng Ye provided guidance on algorithm development and wet lab experiments, respectively. Wenqing Yan provided advice on target selection. Yuhang Chen assisted with algorithm development. All authors reviewed and approved the final manuscript. Junyu Yan and Zibo Cui contributed equally to this work.

## Notes

### Competing Interest Statement

The authors have declared no competing interest.

