## Supplementary information for "Robust and Reliable *de novo* Protein Design: A Flow-Matching-Based Protein Generative Model Achieves Remarkably High Success Rates"

### A Overview

Comprehensive details regarding the algorithm, methodological framework, and experimental protocols are included in the supplementary materials. Appendices B-J are supplementary materials for the paper. File SI_CD contains wet experimental circular dichroism (CD) spectroscopy results. File SI_TM contains wet experimental Thermal Melting results. File SI_SPR experimental contains wet experimental Surface Plasmon Resonance (SPR) File SI_SDS contains wet experimental Sodium Dodecyl Sulfate-Polyacrylamide Gel Electrophoresis (SDS-PAGE) results..

### B Flow matching for generative protein backbone design

#### B.1 Optimal transport flow matching model

Deep generative models are advanced deep learning algorithms for estimating and sampling unknown data distributions. Recent progress has expanded these models into various forms like Diffusion[1, 2], SDE[3], ODE[3–5],and CNF[6], which share close connections and can be transformed into each other. The Flow Matching model is an efficient simulation-free approach to training CNF models and allowing the adoption of general probability paths to supervise CNF training. It focuses on modeling a target vector field to generate the desired probability path, enhancing CNF training efficiency.

Let $x_1$ denote a random variable distributed according to some unknown data distribution $q(x_1)$. We assume we only have access to data samples from $q(x_0)$ but have no access to the density function itself. Furthermore, we let pt be a probability path such that $p_0 = p$ is a simple distribution, e.g., the standard normal distribution $p(x) = \mathbb{N}(x|0, I)$ .

The objective of Flow Matching (FM) is to obtain a vector field $u_t(x)$ that generates the target probability density path $p_t(x)$.

$$\mathcal{L}_{FM}(\theta) = \mathcal{E}_{t,p_t(x)} \|v_t(x) - u_t(x)\|^2. \tag{1}$$

According to CNF, once the optimized vector field $v_t(x)$ approximates $u_t$, the learned CNF model is capable of generating a probability density path $p_t$, which will enable us to flow from $p_0$ to $p_1$. Given the challenge of directly computing $u_t$, Lipman and colleagues introduced the Conditional Flow Matching (CFM) objective[7]:

$$\mathcal{L}_{CFM}(\theta) = \mathcal{E}_{t,p_t(x)} \|v_t(x) - u_t(x|x_0)\|^2. \tag{2}$$

where $t \sim U[0, 1]$, $x_1 \sim q(x_1)$, and now $x \sim p_t(x|x_1)$. Unlike the FM objective, the CFM objective allows for the easy sampling of unbiased estimates as long as we can efficiently sample from $p_t(x|x_0)$ and compute $u_t(x|x_0)$, both of which can be readily achieved since they are defined on a per-sample basis. They have demonstrated that optimizing the CFM objective is equivalent to optimizing the FM objective, but CFM is easier to model.

This representation of the transition from $x_0$ to $x_1$ through $u_t$ can be expressed using an ordinary differential equation (ODE) model on time $t \in [0, 1]$, as follows:

$$\frac{d}{dt}x_t = u(x_t, t)dt. \tag{3}$$

In constructing $u_t$ and $p_t$, an arguably more natural choice for conditional probability paths is to set them to drive the flow to follow the direction $(X_1 - X_0)$ of the linear path pointing from $X_0$ to $X_1$ as much as possible. This can be achieved by:

$$\min_u \int_0^1 \mathbb{E}\left[\|(X_1 - X_0) - u(X_t, t)\|^2\right] dt, \tag{4}$$

$$x_t = tx_1 + (1-t)x_0, \quad x_0 \sim \mathcal{N}(x|0, I) \tag{5}$$

The conditional probability paths can also be interpreted as a Gaussian distribution with linearly varying mean and standard deviation, where $\mu_t(x) = tx_0$, and $\sigma_t(x) = (1-t)$; thus, $\mathcal{N}(x_t; tx_0, (1-t)^2 I)$. The vector field $VF$ can then be defined as:

$$u_t(x|x_1) = \frac{x_1 - x_t}{1-t}$$

Consequently,

$$\frac{dx_t}{dt} = c = x_1 - x_0$$

sampling using Euler's method,

$$x_{t+\delta t} = x_t + u_t \cdot dt$$

The above is a succinct description of the FM approach. Compared to the well-known variance-exploding(VE) and variance-preserving(VP) processes[3], the advantage of the OT process lies in its simpler path, which can significantly accelerate training speed. Its signal-to-noise ratio is primarily determined by time $t$. Similarly to VP and VE processes, OT can also be extended to the forms of SDE and flow ODE, allowing for the application of numerous training and sampling techniques.

#### B.2 The SDE and ODE forms of the OTVF (Optimal Transport Vector Field).

**Constant velocity ODE**: Since the mapping from $x0$ to $x1$ that we seek is a solution to the aforementioned Vector Field (VF) $vt$, it can thus be expressed in the form of an Ordinary Differential Equation (ODE):

$$dx_t = v(x_t, t)dt$$

According to $x_t = tx_1 + (1-t)x_0$, we can derive the target ODE: $\frac{dx_t}{dt} = (x_1 - x_0)dt$. Therefore, the optimization objective is to ensure $v(x_t, t) \approx (x_1 - x_0)$, that is,

$$\hat{\theta} = \arg\min_{\theta} \mathbb{E}_{t,x_t} \left\| x_1 - x_0 - v_\theta(x_t, t) \right\|_2^2 . \tag{6}$$

**SDE**: The expression $x_t = t * x_1 + (1-t) * x_0$ describes $p(x_t|x_0, x_1)$, and its continuous transition process $p(x_{t+\delta t}|x_t)$ can be represented as an SDE.

The aforementioned probability path can be viewed as a special form of a Gaussian noise process. According to the diffusion model, the corresponding noise kernel parameters $(\alpha_t, \sigma_t)$, where $\alpha_t = t$, $\sigma_t^2 = (1-t)^2$,

We can also express its forward SDE, reverse SDE, and flow ODE forms, which can be parameterized through $(f_t,\ g_t)$.

For a general SDE

$$dx = f_t(x)dt + g_t dw$$

, If a $p(x_t|x_1) \sim N(x_t; \alpha_t x_1, \sigma_t^2 I)$

Then,

$$\alpha_{t+\Delta t} = (1 + f_t \Delta t)\alpha_t, \tag{7}$$

$$\sigma_{t+\Delta t}^2 = (1 + f_t \Delta t)\sigma_t^2 + g_t^2 \Delta t \tag{8}$$

Let $\Delta t \sim 0$

$$f_t = \frac{d}{dt}(\ln \alpha_t) = \frac{1}{\alpha_t}\frac{d\alpha_t}{d_t}, g_t^2 = \alpha_t^2 \frac{d}{dt}\left(\frac{\sigma_t^2}{\alpha_t^2}\right) = 2\alpha_t\sigma_t \frac{d}{dt}\left(\frac{\sigma_t}{\alpha_t}\right)$$

Therefore, we obtain the SDE coefficients of OTVF as $f_t = -\frac{1}{t}$ and $g_t = \sqrt{\frac{2}{t} - 2.}$, and forward process:

$$dx = f(x_t, t)dt + g(x, t)dw,$$

$$dx = -\frac{1}{t}x_t dt + \sqrt{\frac{2}{t} - 2dw}$$

Its corresponding reverse SDE is:

$$dx = \left[f_t(x) - g_t^2 \nabla_x \log p_t(x)\right] dt + g_t dw$$

$$\nabla_x \log p_t(x) = -\frac{x_t - tx_1}{(1-t)^2}$$

**Probability Flow ODE**: Considering the reverse process of the aforementioned SDE, we can obtain its Probability Flow ODE solution:

$$dx = \left\{ f(x_t, t) + \frac{g_t^2}{2} \nabla \log p_t \right\} dt,$$

It can also be reformulated as:

$$v_t(x_t, t) = f(x_t, t) + \frac{g_t^2}{2} \nabla \log p_t,$$

The sampling method corresponding to the solution of the Probability Flow ODE is mathematically equivalent to that of the Constant Velocity ODE, yet this form of solution allows for more conditional controls.

#### B.3 Training with Reconstruction Errors.

The model is trained with losses across various dimensions. In Section Protein Structure Initialization, we describe a representation method for protein residues as SO(3) rotation matrices and a translation vector.

**Trans loss**: For the translation part, the loss function can be expressed as the sum of squared errors of the translation vector predictions:

$$L_{\text{trans}} = \sum_{n=1}^{N} \left\| \mathbf{v}_x^{(n)}(T_t, t) - \dot{\mathbf{x}}_t^{(n)} \right\|^2$$

where $\mathbf{v}_x^{(n)}(T_t, t)$ represents the predicted translation vector field for the $n$-th residue at time $t$, and $\dot{\mathbf{x}}_t^{(n)}$ is the calculated translation velocity according to the formula:

$$\dot{\mathbf{x}}_t^{(n)} = \frac{\mathbf{x}_1^{(n)} - \mathbf{x}_t^{(n)}}{1 - t}$$

**Rotations loss**: For the rotation part, the loss function can be expressed as the sum of squared errors of the logarithmic map predictions of the rotations:

$$L_{\text{rot}} = \sum_{n=1}^{N} \left\| \mathbf{v}_r^{(n)}(T_t, t) - \dot{\mathbf{r}}_t^{(n)} \right\|^2$$

where $\mathbf{v}_r^{(n)}(T_t, t)$ represents the predicted rotation vector field for the $n$-th residue at time $t$, and $\dot{\mathbf{r}}_t^{(n)}$ is the calculated rotation velocity according to the formula:

$$\dot{\mathbf{r}}_t^{(n)} = \frac{\log_{\mathbf{r}_0^{(n)}}(\mathbf{r}_1^{(n)})}{1 - t}$$

**Atoms loss**: The backbone atoms loss is a constraint applied to the four backbone atoms. The loss is calculated as follows: the backbone atom loss, *bb_atom_loss*, is calculated by summing the squared differences between the ground truth $\mathbb{X}_1$ and predicted backbone$\mathbf{X}_1$ atoms, weighted by the loss mask. This sum is then normalized by the time denominator:

$$L_{\text{atom\_loss}} = \frac{(\mathbb{X}_1 - \mathbf{X}_1)^2}{\text{1-t}}$$

This loss function enforces a constraint on the prediction of backbone atom positions, aiming to minimize the squared error between predicted and true positions, scaled and normalized appropriately.

**Pairwise distance loss**: represents the error in the predicted pairwise distances between backbone atoms compared to the ground truth distances. It focuses on those atom pairs specified by the loss mask, aiming to minimize discrepancies in the spatial relationships between atoms within protein backbones. This loss helps ensure that the predicted protein structure preserves the geometric properties of the actual protein structure.

First, the predicted and ground truth atomic positions are flattened, resulting in sequences of atom positions with dimensions $[\mathrm{Batch}, \mathrm{num\_res} \times 4]$. Subsequently, the mean squared error (MSE) is calculated between every pair of atomic positions, forming distance matrices of dimensions $[\mathrm{Batch}, \mathrm{num\_res} \times 4, \mathrm{num\_res} \times 4]$. An L2 loss is then computed between the two resulting distance matrices obtained for the ground truth and the predictions.

**Violations loss**: To reduce the clashes in generated structures, a method similar to that of AlphaFold2, known as "structural violations," is employed.

##### B.4 Training with likelihood on an Evidence Lower Bound.

Denoising loss in diffusion models can often be parameterized by a denoising neural network $\hat{x}_\theta(x, t)$ that is trained to predict $x_0$ given a noisy sample $x_t$. This process typically involves minimizing a denoising loss, primarily manifesting as a decrease in reconstruction error. However, our training observations revealed that exclusively utilizing this method yields protein structures of high density, resembling compact globular proteins, with an increase in contact order proportional to the length, and an excess of helices in the structure. Hence, taking inspiration from the Chroma model, we devised a pseudo-ELBO loss.

### C Protein structure representation and forward process

##### C.1 Protein structure representation in SE3 space and linear transformation Euclidean space

The structure of a protein is an ordered three-dimensional point cloud, and the most direct representation method is the sequence of atomic positions in Euclidean space. Compared to common structures like text, images, and point clouds, its characteristics include a large spatial span and high structural precision. For a protein of length 100, its radius of gyration often falls around 60Å ($R_g \propto rN^\nu, \nu \sim 0.4$)[8, 9], but the atomic positions of proteins are strictly constrained by physical and chemical rules. For subsequent protein design tasks, if the structural perturbation is around 0.2Å, it would significantly increase the difficulty of structural design.[10, 11] This also leads to the challenge of designing long-sequence proteins, namely the extrapolation problem of protein structure generation models.

If diffusion is directly applied to the original Cartesian coordinates, then a significant amount of computational power would be wasted on basic rules such as chains and densities. Moreover, it would severely limit its extrapolation and accuracy, with the model performance rapidly decreasing as length increases[12]. Since the positions of different atoms on the same amino acid are not independently and identically distributed, diffusion in the $\mathbb{R}^3$ space either involves only diffusing the $C_\alpha$ atom and representing the remaining atoms on the same amino acid as functions of the $C_\alpha$ atom[13], significantly reducing computational complexity. Some attempts have been made to perform diffusion in SE3 space, where in Cartesian space, diffusion is only needed for the $C_\alpha$ atoms, and the remaining three atoms are represented by rotation matrices[14]. Through the efforts of some researchers, the diffusion model generation in Riemannian space has been developed[15]. However, during the generation process, there are some practical challenges in synchronously handling both SO3 and Cartesian spaces[16], especially when conducting conditional generation and multiple sampling techniques, such as the issue of conditional derivation in SO3.

**Represented as an SE3 frame**: Similar to AlpahFold2[17], it uses the positions of a few basic backbone atoms to calculate an orthogonal matrix and represents translation with the position of the $C_\alpha$ atom. The calculation method is as follows:

$$\vec{X}_{bb} = [\vec{X}_N, \vec{X}_{C_\alpha}, \vec{X}_C, \vec{X}_O]$$

$$\vec{u}_{C_\alpha - N} = \frac{\vec{X}_N - \vec{X}_{C_\alpha}}{\sqrt{\left|\vec{X}_N - \vec{X}_{C_\alpha}\right|^2 + \epsilon}} \; \vec{u}_{C_\alpha - C} = \frac{\vec{X}_C - \vec{X}_{C_\alpha}}{\sqrt{\left|\vec{X}_C - \vec{X}_{C_\alpha}\right|^2 + \epsilon}}$$

$$\vec{n}_1 = \vec{u}_{C_\alpha - N} \; \vec{n}_2 = \frac{\vec{n}_1 \times \vec{u}_{C_\alpha - C}}{\sqrt{\left|\vec{n}_1 \times \vec{u}_{C_\alpha - C}\right|^2 + \epsilon}} \; \vec{n}_3 = \frac{\vec{n}_1 \times \vec{n}_2}{\sqrt{\left|\vec{n}_1 \times \vec{n}_2\right|^2 + \epsilon}}$$

$$\vec{r} = \begin{bmatrix} \vec{n}_1 & \vec{n}_2 & \vec{n}_3 \end{bmatrix}$$

$$\vec{T} = C_\alpha$$

$$\vec{X}_{bb} = [\vec{r}, \vec{T}]$$

**Represented as an $\mathbb{R}^3$ frame**: Similar to Chroma[12], We represent the three atoms N, C, O of the protein sequence structure $X$ as the difference relative to the $\mathrm{C}_\alpha$ atom. Such a structure can be represented by a linear transformation matrix $R$:

$$\vec{X}_{bb} = Rz$$

$$R_{\text{residue}} = \begin{bmatrix} \sigma_{\mathrm{N}} & 0 & 0 & 0 \\ 0 & \sigma_{C\alpha} & 0 & 0 \\ 0 & \sigma_{C\alpha} & \sigma_{\mathrm{C}} & 0 \\ 0 & \sigma_{C\alpha} & 0 & \sigma_{\mathrm{O}} \end{bmatrix}$$

The transformation $T$ from the $z$ space to the $X$ space can be expressed as a linear transformation matrix. Where $\sigma_{C_\alpha} = R_g, \sigma_{N,C,O} = 1, z \in \mathbb{N}(0, 1)$

**Symmetries**: Prior research [18, 19] has highlighted the critical role of integrating the target system's symmetries into the flow model. We normalize the overall protein structure using the zero center of mass (CoM) subspace of $\mathbb{R}^N \times 3$, which also helps to reduce the distance variations caused by rotation. We also performed an alignment between the randomly sampled noise structures and the real structures. According to Klein et al[20]., this method can significantly reduce the increase in kinetic energy caused by rotation and greatly enhance training efficiency while reducing the transport cost during the sampling process.

$$COM \;\; SE3: \;\; T = T - mean(C\alpha)$$

$$COM\ X:\ \ X = X - mean(C_\alpha)$$

#### C.2 Protein Structure Initialization

At the beginning of the forward process, for any given data $X_1$, we initialize a corresponding random structure $X_0$. The specific sampling and processing methods are as described in Protein Initialization. For this random structure, we process it according to the aforementioned symmetry requirements and align it with the $X_1$ structure. After adjustment, we obtain a randomly sampled structure $\vec{X_{bb}} \in \mathbb{R}^{N\times 3}$ represented in the $\mathbb{R}^{N\times 3}$ space.

**Algorithm 1** Protein Initialization

**Function** Protein Initialization$(X, N, R, Z, \sigma_{C_\alpha}, \sigma_{\mathrm{N,C,O}})$
**for all** $l \in [1, \ldots, N]$ **do**
    $X \in \mathbb{R}^{N\times 3}$ ▷ 3D point cloud of protein structure
    $X = RZ$ ▷ Apply linear transformation
    $\sigma_{C_\alpha} = R_g$, $\sigma_{\mathrm{intra}} = 1$ ▷ Set normalization parameters
    Normalize $X$ using zero CoM ▷ Symmetry handling
    Pre-align with Kabsch ▷ Align prior and data
    $Z_{C_\alpha} \sim \mathcal{N}(0, 1)$ ▷ Sample backbone
    Optimal align $\tilde{Z_{C_\alpha}}$ with $X_{C_\alpha}$ ▷ Alignment
    $Z_N, Z_C, Z_O \sim \mathcal{N}(0, 1)$ ▷ Sample side chains
    $Z = [Z_N, Z_{C_\alpha}, Z_C, Z_O]$ ▷ Form complete sample
**end for**
**Return** $X, Z$ ▷ Apply OT Flow Matching for training

#### C.3 Forward Process

rotein structures can be parameterized using two methods: $\mathbb{SE}^3$ and $\mathbb{R}^3$. When perturbing the original structure during the forward process, operations can be conducted in either $\mathbb{SE}^3$ or $\mathbb{R}^3$. space. The sensitivity of the perturbed structures to time varies between these two spaces.

##### C.3.1 Noising in SE3 space

After obtaining the initial prior distribution of random structures, perturbations can be applied synchronously in both R3 and SO3 spaces with reference to [16, 21]. For translations in the R3 space, the perturbation can be expressed as

$$\text{Translations}(\mathbb{R}^3): \vec{T} = (1-t)\vec{T_0} + t\vec{T_1}$$

For rotation matrices in the SO(3) space, where a random sample $\vec{X_0}$ is drawn from a uniform distribution, to obtain the optimal transport path to the target sample $\vec{X_1}$, one can use the geodesic approach, following Yim et al.[21]. This allows us to define

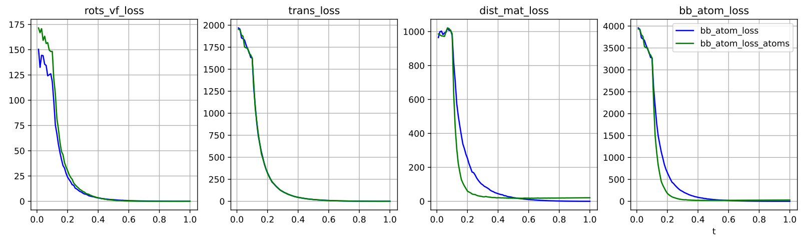


**Fig. 1**: The variation of loss over time for different perturbation methods. The blue curve represents the structural difference between the original structure and the structure after noise addition in the SE(3) space, while the green curve illustrates the difference after noise addition in the $\mathbb{R}^3$ space. The vertical axis quantifies these differences using loss values, calculated using the same methodology employed during training. Overall, adding noise in the SE(3) space results in more gradual structural changes.

$$\text{Rotations}(\text{SO}(3)) : \vec{r}_t = \exp(\vec{r}_0 t \log(\vec{r}_0^{-1} \vec{r}_1))$$

In the $\mathbb{R}^3$ space, since all positions are atomic, direct superposition of positions is possible:

$$\vec{X}_t = (1-t)\vec{X}_0 + t\vec{X}_1$$

### D Architecture

#### D.1 The SE3 Frame as an intermediate state in the iteration

Originflow constructs a network that combines an IPA transformer and a GNN. The IPA transformer[17] is a structural prediction module widely used in the protein domain, but the performance of the transformer structure is significantly influenced by length (i.e., the extrapolation problem of transformers[22]). Therefore, after obtaining the initial structure from the transformer, we employ a local GNN network to refine the positions of protein atoms.

During the denoising process, the protein structure is represented in SE3. The model initially extracts node features and pair features, which are then fed into the IPA transformer to predict a preliminary denoised structure. Subsequently, for the $C_\alpha$ atoms, a K-nearest neighbors approach is used to find the surrounding K amino acids, constructing a graph. Then, the corresponding K nearest neighbor features from the pair features are extracted to form the new edge features for the GNN.

We conducted comparative experiments between IPA only and the overlay of GNN methods. When using the scope dataset and training for 50k steps, the model employing the GNN structure demonstrated better sampling quality.

**Represented as an $\mathbb{R}^3$ frame**: Similar to Chroma[12], We represent the three atoms N, C, O of the protein sequence structure $X$ as the difference relative to the $C_\alpha$ atom. Such a structure can be represented by a linear transformation matrix $R$:

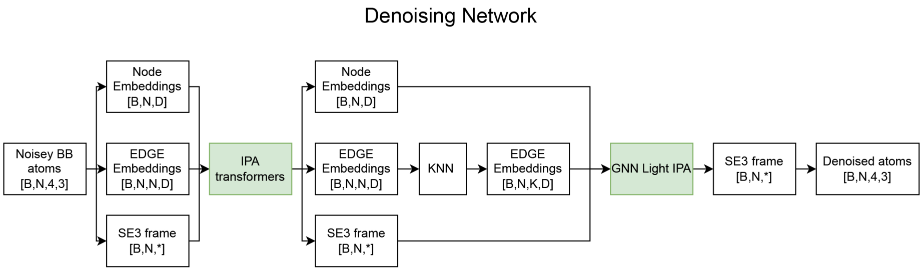


**Fig. 2**: Overview of the **Originflow** architecture, which integrates a global Transformer module with a local GNN for protein structure generation. This hybrid design enables effective modeling of both long-range dependencies and local structural constraints.

**Features**: The computation of input features for the model is as follows features. Compared to basic features such as position or time embedding, and composite basic features like *cross-node*, *cross-position*, and *distogram*, the **Internal Features** geometric structural features significantly accelerate convergence speed, reduce loss, and yield better quality during sampling.

| Node features | |
|---|---|
| Internal Features [1, L, 10] | The protein backbone chain -[N]-[CA]-[C]- bond lengths (NCaC_L), bond angles (NCaC_A), and dihedral angles (NCaC_D) were calculated; as well as the bond lengths (O_L), bond angles (O_A), and dihedral angles (O_D) for the peptide bond [C]=O. The sine and cosine values of the angles and dihedrals were computed, and the logarithm of the bond lengths was taken. Subsequently, the aforementioned features were combined. |
| Pos embedding [1, L, 128] | Cosine Positional Encoding. |
| time embedding [1, L, 128] | Embedding of time step. |
| Pair distance rbf [1, L, 16] | Calculate the relative distances between atoms for different pairs (i, j), and compute the RBF (Radial Basis Function) mapped distances. |
| Pair distance chain [1, L, 3] | Contains three features, namely whether they are on the same chain, the sequence number distance on the chain, and the sign of the intra-chain distance. |
| Cross node [1, L, 128] | Perform cross concatenation of the obtained node features, i.e., $cross_{i,j} = concat(node_i, node_j)$, and then map it through a linear layer. |
| Cross pos [1, L, 128] | Subtract the chain distances of two positions and map through an embedding. |
| $C_\alpha$ distogram [1, L, 22] | Calculate the distogram using $C_\alpha$ atoms, with $min_{bin} = 1e-3$, $max_{bin} = 20.0$, and $numberofbins = 22$. |
| $sc$ $C_\alpha$ distogram[1, L, 22] | Calculate the distogram using $C_\alpha$ atoms of self condition data, with $min_{bin} = 1e-3$, $max_{bin} = 20.0$, and $numberofbins = 22$. |

In terms of features, for binder design, it is not only important to consider the backbone structure but also the amino acid sequence and the manner of side-chain packing. These factors have a significant impact on the protein-protein interface. Hence, we have introduced some additional features in the binder design mode.

| Node features | |
|---|---|
| AA type Embedding [1, L, 21] | 20 amino acids and one missing value X are mapped to 21-dimensional features. |
| Physicochemical embedding [1, L, 571] | 571 types of physicochemical features for each amino acid are extracted, with the missing value features being the average of other amino acids' features. |
| Chi embedding [1, L, 8] | Up to four side-chain dihedral angles, with both sin and cos values computed for each. |
| B factor embedding [1, L, 128] | The B factor is encoded into four mapping sequences using bins thresholds of 15, 30, 45, 60, along with the original B factor, then combined and mapped to 128-dimensional features through a fully connected layer. |

#### D.2 GNN Used for Local Structure Optimization

The GNN utilizes initial structural features combined with the output of IPA to construct node and edge features. On each layer of the graph, after updating node features using adjacent nodes.

We designed three update mechanisms: FFN, nodelightIPA, and neighlightIPA. The feature processing and edge-node information update strategies for these modules are consistent. The difference lies in how the updated edge-node information is utilized to update the structure.

- **FFN**: The updated edge information is directly passed through a PositionWiseFeedForward function to produce the output vector for structural correction.
- **nodelightIPA**: The updated node features, serving as the single feature input in IPA, with the pair feature being the Node, and concurrently inputting the current structure, are updated through the IPA transformer.
- **neighlightIPA**: According to the knn graph topology, the node features of neighboring nodes $[B, N, D]$ are aggregated into $neigh\_feature[B, N, K, D]$, and adjusted to $[B{*}N, K, D]$. Within the $K$ neighboring nodes, the lightIPA processing is applied, thereby achieving updates that incorporate neighborhood information within each neighborhood. At a position $i$ of an amino acid, which may appear in the neighborhood of $M$ points, it will be updated $m$ times. We fuse the outputs of these $m$ IPA processes, *IPAEmbed*, and then pass it through an update layer, outputting the correction values for rotation and translation to update the structure.

$$F_{neigh} = gather \quad nodes(nodes, edge\_idx)$$

$$\underset{[b*n,k,d]}{F_{\text{neigh}}} \leftarrow \underset{[b,n,k,d]}{F_{\text{neigh}}}$$

$$\underset{[b*n,k,d]}{F^{\text{update}}_{\text{neigh}}} = LightIPA(F_{\text{neigh}}, R)$$

- $F_{\text{neigh}} \in \mathbb{R}^{B\times N\times K\times D}$ is the feature matrix for neighbor nodes.
- $I_{\text{neigh}} \in \mathbb{R}^{B\times N\times K}$ is the index matrix for neighbor nodes.

Given: For each node $i$, we aggregate the features $F^{\text{update}}_{\text{neigh}}$ corresponding to all occurrences of $i$ in $I_{\text{neigh}}$, and assign it to a new feature tensor $F_{\text{node}}$:

$$F_{\text{node},i} = \bigoplus_{(b,j,k)\,:\,I_{\text{neigh},b,j,k}=i} F^{\text{update}}_{\text{neigh},b,j,k}$$

where:

- $F_{\text{node},i}$ represents the new feature for node $i$.
- $\bigoplus$ denotes an aggregation operation, which can be sum, mean, or any appropriate aggregation function.
- $I_{\text{neigh},b,j,k} = i$ signifies that in batch $b$, within the neighbor list of node $j$, at the $k$-th position, the index of the neighbor node is $i$.

**Algorithm 2** GNN update layer

**Function**Forward$h_V, h_E, E_{\text{idx}}, \text{mask}_V, \text{mask}_{\text{attend}}, R, \text{pos}, \text{kwargs}$
$h_{EV} \leftarrow \text{cat\_nodes}(h_V, h_E, E_{\text{idx}})$
$h_{\text{message}} \leftarrow FFN(h_{EV})$
$dh \leftarrow \text{sum}(h_{\text{message}}, -2)/\text{scale}$
$h_V \leftarrow \text{norm}_1(h_V + \text{dropout}_1(dh))$
$dh \leftarrow \text{dense}(h_V)$
$h_V \leftarrow \text{norm}_2(h_V + \text{dropout}_2(dh))$
$h_{EV} \leftarrow \text{cat\_nodes}(h_V, h_E, E_{\text{idx}})$
$h_{\text{message}} \leftarrow update(h_{EV})$
$h_E \leftarrow \text{norm}_3(h_E + \text{dropout}_3(h_{\text{message}}))$
$h_V, R \leftarrow \text{node\_update}(h_V, R, )$
$h_V \leftarrow h_V + h_{V\_update}$
**Return** $h_V, h_E, R$

#### D.3 Extrapolation

The attention component naturally faces challenges in extrapolation generalization[23], and due to constraints in computational resources, the sequence length during training is often shorter than during inference. For the attention mechanism, we have utilized rotational positional encoding[24] to capture the relative information between different positions. Furthermore, we have scaled the attention scores by a length scale factor to prevent the increase in attention entropy caused by excessive length.

### E Training

#### E.1 Dataset

During our training process, we mainly utilize data from the Protein Data Bank. For training the base model, we select protein structures with a resolution of 2.6Å or less, taking proteins of lengths between 40 and 2048, and cropping them to a length of 384. The protein data includes both homomers and heteromers but excludes small molecules.

For motifs, we also select data with a resolution less than 2.6Å and filter using a redundancy level of 30%. From each cluster, we select one protein, primarily using homomer sequences.

Similarly, for the binder model, we use a resolution of less than 2.6Å and filter with a redundancy level of 30%, but we divide the proteins into protein-protein complexes and non-complexes. For protein-protein (PP) complexes, when the length is less than 450, we take the entire complex. For lengths greater than 450, we select the area near the interface, centering on the point of minimum distance, with each side of the complex not exceeding 450 adjacent residues. This selection method may result in discontinuous residues. Therefore, we also include less than 450-length non-complex multichain proteins, totaling 5000 entries. The proportion of data among these three categories is approximately 1:2:3.

#### E.2 Base model and Modifications

The base model is trained using the aforementioned perturbation strategies, data, along with Reconstruction Errors and ELBO.

In the training of the motif and binder design models, we adjusted the structure of the GNN part and the features to accommodate different tasks.We first established an iterative loop, where in each iteration, we use the most recently updated structure (updated through global IPA updates or the GNN updates from the previous cycle) to recompute the nearest neighbors KNN and recalculate the features.

**Motif model**: In the motif model, the fixed mask for the motif region is set to 1, while the remaining positions are set to 0, and the time embedding is calculated using a fixed time of 0 (endpoint time). Adjustments are mainly made in the iteration of the GNN. When obtaining the initial structure using IPA, the fixed part (motif region) does not participate in updates, thus preserving the original structure of the motif region, whereas the scaffold part outside of the motif is generated. Within the GNN network, node and pair features are recalculated and summed with the results from IPA, followed by each GNN update computation. After the GNN performs a neigh update, all positions are updated. However, due to the setting of the time embedding, the deformation in the motif region is minimal.

**Binder model**: We fine-tuned the model using two datasets: PDB complex and PDBBind. Initially, we filtered the complexes with lengths below 2048 residues and applied cluster30 to remove redundancy. For each complex, we calculated the distances between residues of the internal subunits. We extracted the closest pairs of residues, not exceeding a total of 384 amino acids, as the interface data. Using this data, we fine-tuned the model for 10 epochs.

Subsequently, the Binder model was fine-tuned using 2000 protein-protein complexes from the PDBBind database. The data processing method was identical to the one described above. The feature processing in the Binder model differs from the base and motif models. Specifically, we enhanced the sequence portion: for the target chain, its sequence was mapped to a 128-dimensional feature vector, which was then merged with other node features.

The most significant difference between the Binder model and the Motif model lies in the recalculation of the GNN topology at each GNN encoder layer, and the fixed area in the GNN part is not updated. This is because binder design is more sensitive to structural changes. The Binder model also predicts the b-factor, side-chain Chi angles, and amino acid sequence at each position.

**Fine-tuning in Training Precision:** Initially, we trained the base model for 50 epochs using 16-bit mixed precision at a sequence length of 384. Subsequently, we fine-tuned the model for 5 epochs using 32-bit precision at a sequence length of 320. Both the motif and binder models were fine-tuned using 16-bit mixed precision. We tested the generative capabilities of models trained with different precisions in the IN SICILI test section.

#### E.3 Sampling

**Overall Sampling Method:** Our overall sampling method is based on the ODE Euler method, but it manifests in different forms.

**In CV ODE:** The sampling method used is as follows:

$$\hat{X} = Denoiser(X_t, t)$$

$$\frac{dx}{dt} = v = \frac{\hat{X} - X_t}{1 - t}$$

$$X = X_t + v * dt$$

Specifically,

$$X_t = (r_t, trans_t), \hat{X}_1 = (\hat{r_1}, tr\hat{a}ns_1)$$
$$trans_{t+1} = trans_t + \frac{\hat{t}rans_1 - trans_t}{1 - t}$$
$$r_{t+1} = exp_{rt}((1 - e^{-ct} log_{rt}(\hat{r_1})))$$

**In PF ODE**: The sampling method used is as follows:

$$h = \frac{1}{t}, g^2 = 2 * \frac{1 - t}{t}$$

$$\nabla \log p = \frac{t * tr\hat{a}ns_1 - trans_t}{1 - t}^2$$

$$vf = h * \hat{trans_t} + \frac{g^2}{2} * \nabla \log p * \lambda$$
$$trans_{t+1} = trans_t + vf * d_t$$

**Mathematical Equivalence:** The two methods are mathematically equivalent; however, the solution approach in PF ODE is more conducive to introducing control variables.

**Unconditional and Conditional Generation:** For unconditional generation, we primarily utilize the generation method of CV ODE, while for conditional generation, we predominantly employ the generation method of PF ODE.

**Sequence Generation:** For the generated structures, we design sequences for the generated backbone using the ProteinMPNN Ca only mode, with a weight of $v\_48\_020$, and a sampling temperature set to the default value of 0.1.

### F In silico experiment

#### F.1 Unconditional benchmarking

**Backbone Generation**

For lengths ranging from 40 to 600, incrementing by 10, we use the base model trained with 16-bit mixed precision to generate 10 backbones, resulting in a total of 560 original structures. We first randomly selected 100 cases and designed different numbers of sequences using MPNN, followed by folding with ESMFold. We found that after designing four sequences, the average pLDDT stabilized above 80, and RMSD was below 3Å. The marginal improvement diminished after designing eight sequences. Therefore, for all subsequent experiments, we designed eight sequences for each structure and selected the one with the smallest RMSD for further calculations.

**Model Parameter Selection** We first trained the base model using FP16 mixed precision, with a maximum training sequence length of 384. We then fine-tuned the model for five epochs using FP32, but with a maximum length of 256. We compared the performance of these two models, and the results showed that the FP16 mixed precision model was more stable and performed better in all aspects . Therefore, all subsequent model generations used the FP16 mixed precision trained model.

**Refolding analysis**

Our TMscore is 0.83, with a median of 0.9, and the RMSD mean is 3.22, with a median of 1.92. This indicates that the structures generated by our model possess high designability. Additionally, the slope is nearly zero and the correlation coefficient is around 0.2, indicating that the model exhibits excellent generalization across different lengths, suggesting that the model's capability does not significantly deteriorate with increasing length. The mean and median pLDDT values are 77.5 and 80.83, respectively, which are relatively high, demonstrating good stability along with pAE, considering the inherent length attenuation in both ProteinMPNN and the structural prediction model.

**Geometry statistics benchmark**

We compared the structural generation results of different models. Specifically, we compared monomer generation using Chroma and RFdiffusion, and complex generation using Chroma (RFdiffusion requires a larger GPU for complex calculations).

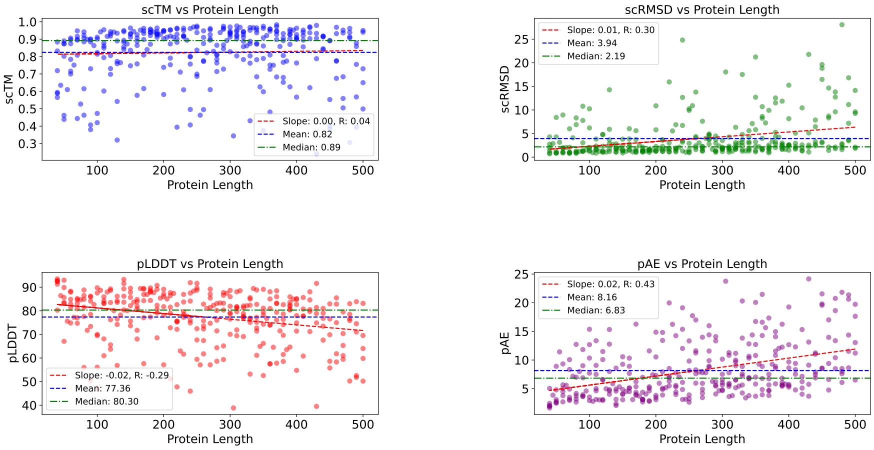


**Fig. 3**: **Refolding scores of monomer structures generated by Originflow across different lengths.**
Due to constraints in hardware memory and prediction duration, we conducted refolding experiments on structures shorter than 500. **Slope** and **R** refer to the fitting slope and correlation coefficient of scores relative to length, respectively. **Mean** and **Median** refer to the average and median values, respectively.

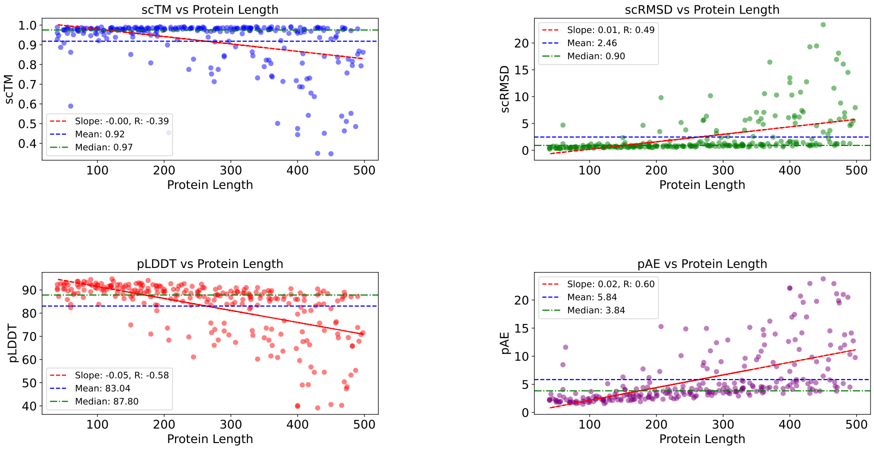


**Fig. 4**: **Metrics of refolded monomer structures generated by RFdiffusion relative to length**
After generating structures with RFdiffusion, eight sequences were designed for each structure and sent to ESMFold. The metrics scTM, scRMSD, pLDDT, and pAE were then recorded.

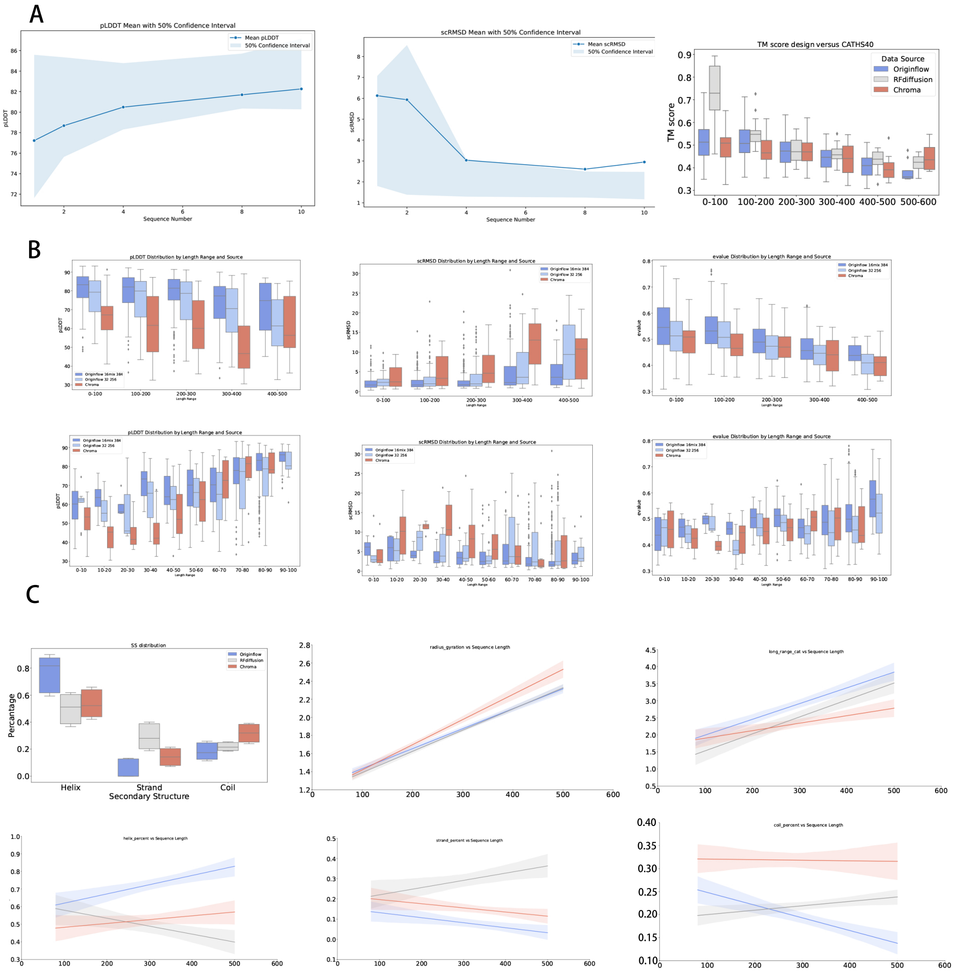


**Fig. 5**: **Various Computational Metrics for Originflow ModelGenerated Structures A.** Performance of pLDDT and RMSD for 100 structures of lengths ranging from 40 to 400, each with varying numbers of sequences filled using ProteinMPNN and re-folded using ESMFold. As the number of sequences increases, pLDDT consistently improves while RMSD shows a decreasing trend. After 8 sequences, the improvement in metrics slows down. **B.** Comparison of monomer generation capabilities among the 16-bit mixed-precision model, the FP32 fine-tuned model, and the Chroma model. The 16-bit mixed-precision model outperforms the FP32 fine-tuned model. **C.** Statistical analysis of the structures generated by Originflow, RFdiffusion, and Chroma models for 300 cases of varying lengths, illustrating the relationship between sequence length and the generated protein structures.

For both monomer and multimer generation, we calculated the following features:

1. Secondary structure using DSSP
2. Radius of gyration
3. Long-range residue connectivity
4. Changes in protein secondary structure with increasing length

The figures illustrate these features.

Firstly, for monomer generation, Originflow tends to focus more on alpha structures, with the least on coil structures. Additionally, in terms of radius of gyration and long-range residue connectivity, Originflow shows an increase similar to RFdiffusion as sequence length increases. A notable characteristic of our model in secondary structure is that the coil decreases significantly with increasing sequence length, while alpha structures increase substantially. This indicates that with longer sequences, the model enhances overall stability by increasing the proportion of more stable alpha structures.

**Diversity analysis**

To evaluate the diversity of the structures generated by our model, we calculated the pairwise TM-scores (taking the maximum value) of the structures generated by the same model. We compared these results with those of Chroma and RFdiffusion. Our TM-scores (mean: 0.45, median: 0.44) are significantly lower than those of Chroma (mean: 0.56, median: 0.54) and RFdiffusion (mean: 0.48, median: 0.47).

Additionally, we performed multidimensional scaling (MDS) projection based on the pairwise similarities, mapping and clustering the structures accordingly. The results show that the clusters of Chroma are more concentrated, whereas our model's structures are more evenly distributed. This indicates that our generated structures exhibit significantly higher diversity.

**Novelty** We used Foldseek to compare the generated structures with the CATH 4.3 S40 structures based on similarity, employing the TM Score to assess the closeness of the generated structures to native structures. For each generated sample, the PDB structure with the highest TM Score hit identified by Foldseek was considered as the similar structure. We utilized the overall distribution of TM Scores among the generated samples as an indicator of innovativeness, as illustrated in Fig.

We conducted a comparative analysis between the samples generated by RFdiffusion, Chroma, and our model. It was observed that Chroma exhibits the least similarity with the native dataset(mean aln tmscore 0.59), whereas RFdiffusion shows the greatest(mean aln tmscore 0.66), with our samples positioned in between(mean aln tmscore 0.61). This finding is corroborated by the refolding experiments mentioned above, indicating that Chroma possesses the greatest structural innovativeness. In contrast, sequence generation models like ProteinMPNN, which rely on the distribution of the training data, often show poorer performance in generating innovative structures and in refolding aspects.

| | Originflow | Evodiff | RF diffusion |
|---|---|---|---|
| Successed Cases | 16 | 12 | 13 |
| Media Success Rate(%) | 13 | 12 | 25 |

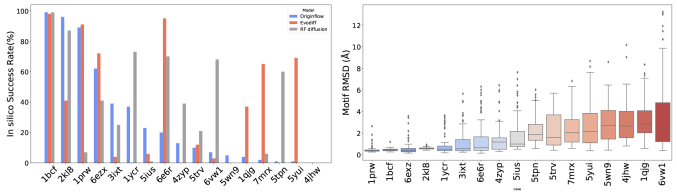


**Fig. 6**: The performance of Originflow in solving the motif-scaffolding problems

### G Conditional design for Motif

Motif scaffolding is a critical component in the de novo design of proteins. We constructed a dataset comprising 17 motifs, drawing inspiration from RFDiffusion and PVQD. Each motif in the dataset was published after the cutoff date for the training data. The dataset includes various types of motifs such as viral epitopes, receptor traps, ligand interfaces, small molecule binding sites, and enzyme active sites.

For each motif, we generated 100 structures. We used ProteinMPNN to fill in the sequences, which were then subjected to structure prediction using ESMFold. A generated structure was considered a successful solution to the motif design problem if its overall pLDDT score was greater than 70 and its motif RMSD was less than 1.0 Å.

Our model successfully solved 16 out of the 17 motif design problems, outperforming Evodiff (12 solved), and RFdiffusion (13 solved). SFig.motif

Similar to RFdiffusion, when performing motif-scaffolding with varying lengths of the sequences, our model can successfully solve 24 out of 25 cases. STable:motif

Additionally, our model demonstrated strong performance on the multi-chain motif 6vw1.

### H Conditional Design for Symmetric Structures

Polyhedral symmetric structures are common macromolecular geometries that play important roles in protein stability and functional efficiency. We adopted a method similar to RFdiffusion, applying symmetric matrices to generate symmetric structures. Without the need for fine-tuning, we can superimpose symmetric rotational operations on the base model, motif model, and SS fine-tuning model to generate symmetric structures. During structure generation, the initially generated SE3 structure ($X_1^{(SE3)}$) is transformed into atomic coordinates within the complete R3 framework ($X_1^{(R3)}$). Based on the type of symmetry, a rotation matrix is added to the divided subunits to obtain ($X_1^{sym(R3)}$), which is then mapped back to ($X_1^{sym(SE3)}$). This mapping guides the iterative steps to generate ($X_{t-1}^{sym(SE3)}$).

| Name | Description | Input | Successed Cases |
|---|---|---|---|
| 1BCF | Di-iron binding motif | 8-15,A92-99,16-30,A123-130,16-30,A47-54,16-30,A18-25,8-15 | 99 |
| 2KL8 | De novo designed protein | A1-7,20-20,A28-79 | 96 |
| 1PRW | Double EF-hand motif | 5-20,A16-35,10-25,A52-71,5-20 | 89 |
| 6EXZ_long | RNA export factor | 0-95,A28-42,0-95 | 67 |
| 6EXZ_med | RNA export factor | 0-65,A28-42,0-65 | 62 |
| 6EXZ_short | RNA export factor | 0-35,A28-42,0-35 | 57 |
| 3IXT | RSV F-protein Site II | 10-40,P254-277,10-40 | 39 |
| 1YCR | P53 helix that binds to Mdm2 | 10-40,B19-27,10-40 | 37 |
| 5IUS | PD-L1 binding interface on PD-1 | 0-30,A119-140,15-40,A63-82,0-30 | 23 |
| 6E6R_long | Ferridoxin Protein | 0-95,A23-35,0-95 | 20 |
| 5TRV_short | De novo designed protein | 0-35,A45-65,0-35 | 19 |
| 5TRV_long | De novo designed protein | 0-95,A45-65,0-95 | 17 |
| 4ZYP | RSV F-protein Site 4 | 10-40,A422-436,10-40 | 11 |
| 5TRV_med | De novo designed protein | 0-65,A45-65,0-65 | 10 |
| 6E6R_med | Ferridoxin Protein | 0-65,A23-35,0-65 | 9 |
| 7MRX_128 | Barnase ribonuclease inhibitor | 0-122,B25-46,0-122 | 9 |
| 6VW1 | ACE2 interface binding SARS-CoV-2 | E400-510/20-45,A24-42,4-10,A64-82,0-5 | 7 |
| 6E6R_short | Ferridoxin Protein | 0-35,A23-35,0-35 | 6 |
| 5WN9 | RSV G-protein 2D10 site | 10-40,A170-189,10-40 | 5 |
| 1QJG | Delta5-3-ketosteroid isomerase active site | 10-20,A38-38,15-30,A14-14,15-30,A99-99,10-20 | 4 |
| 7MRX_85 | Barnase ribonuclease inhibitor | 0-63,B25-46,0-63 | 4 |
| 7MRX_60 | Barnase ribonuclease inhibitor | 0-38,B25-46,0-38 | 2 |
| 5TPN | RSV F-protein Site V | 10-40,A163-181,10-40 | 1 |
| 5YUI | Carbonic anhydrase active site | 5-30,A93-97,5-20,A118-120,10-35,A198-200,10-30 | 1 |
| 4JHW | RSV F-protein Site 0 | 10-25,F196-212,15-30,F63-69,10-25 | 0 |

**Table 1**: Motif Scaffolding benchmark results

### I Conditional Design for SS Condition

| CATH Classes | CATH Architectures | CATH ID | Domain | SS F1 Score |
|---|---|---|---|---|
| Mainly Alpha | Up-down Bundle | 1.20 | 2ynzC01 | 96.55% |
| | Orthogonal Bundle | 1.10 | 1xmkA00 | 92.64% |
| | Alpha solenoid | 1.40 | 1okcA00 | 93.32% |
| | Alpha/alpha barrel | 1.50 | 3iisM00 | 93.51% |
| | Alpha Horseshoe | 1.25 | 3ro3A00 | 93.79% |
| Mainly Beta | Roll | 2.30 | 2o9sA00 | 77.91% |
| | 3-layer Sandwich | 2.102 | 2bmoA02 | 79.50% |
| | Single Sheet | 2.20 | 6d9nB01 | 81.58% |
| | 3 Propeller | 2.105 | 1n7vA01 | 81.26% |
| | Beta Complex | 2.170 | 3f9xD00 | 82.10% |
| | Distorted Sandwich | 2.70 | 1rwhA02 | 81.59% |
| | Beta Barrel | 2.40 | 1x54A01 | 81.37% |
| | Clam | 2.50 | 4mxtA00 | 81.43% |
| | Sandwich | 2.60 | 4unuA00 | 81.15% |
| | Shell | 2.180 | 5kisB00 | 81.14% |
| alphabeta | Box | 3.70 | 1ud9A00 | 79.75% |
| | 2-Layer Sandwich | 3.30 | 6n3dA01 | 82.22% |
| | Alpha-Beta Complex | 3.90 | 2w8tA01 | 84.01% |
| | 3-Layer(aba) Sandwich | 3.40 | 1hdoA00 | 84.87% |
| | Roll | 3.10 | 3goeA00 | 84.29% |
| | Alpha-Beta Horseshoe | 3.80 | 4fs7A00 | 83.27% |
| | Alpha-beta prism | 3.65 | 5u4hA02 | 83.31% |
| | Alpha-Beta Barrel | 3.20 | 2vxnA00 | 83.95% |
| | Super Roll | 3.15 | 1ewfA01 | 83.90% |
| | 3-Layer(bab) Sandwich | 3.55 | 4jtmA00 | 83.53% |
| | 4-Layer Sandwich | 3.60 | 5xp6A00 | 83.63% |
| | 3-Layer(bba) Sandwich | 3.50 | 1lqtB02 | 83.75% |

**Table 2**: Conditional generation results under secondary structure constraints

In de novo generation tasks, it is sometimes necessary to impose prior structural constraints on specific regions or modify the structure of certain areas[14]. The primary structural constraint is the protein secondary structure (contiguous regions of alpha helix, beta strand, or loop). We fine-tuned the model by introducing secondary structure information annotation features. We first used DSSP to calculate the protein's secondary structure, which was input as a point feature. During training, 50% of the time, we masked 50% of the secondary structure information. During generation, the desired regions were specified with secondary structures, while other regions were filled with zeros.

We conducted several tests using representative PDB secondary structures of CATH Alpha, Beta, and AlphaBeta Architectures as outputs to generate structures. For the generated structures, we compared their secondary structures with the native secondary structures. For the main alpha architectures, the average secondary structure overlap was over 95%. For the main beta architectures, the overlap ranged from 70% to 90%, since pure beta architectures cannot form stable supports, leading to slight misalignments in some coil regions. AlphaBeta architectures showed an overlap of 75% to 85%. We also performed specific secondary structure constraint designs for the TIM-Barrel structure, achieving an overlap consistency of 88% to 92%.

### J Conditional design for Binder

| Target | Reference PDB | Design No. | iPTM | pTM | Length | Secondary Structure |
|---|---|---|---|---|---|---|
| COVID | 7rby | 1 | 0.89 | 0.87 | 20 | 2 helices |
| | | 2 | 0.71 | 0.80 | 90 | 7 $\beta$-strands |
| | | 3 | 0.82 | 0.86 | 72 | 3 helices |
| | | 4 | 0.77 | 0.83 | 142 | 6 helices |
| | | 5 | 0.76 | 0.83 | 95 | 4 helices |
| G12D | 7rpz | 1 | 0.78 | 0.86 | 74 | 4 helices |
| | | 2 | 0.80 | 0.85 | 81 | 3 helices of variable length |
| | | 3 | 0.74 | 0.84 | 94 | 3 helices of variable length |
| | | 4 | 0.83 | 0.85 | 51 | 4 helices |
| GP120 | 5cd5 | 1 | 0.88 | 0.87 | 35 | Double helix |
| | 2ny3 | 1 | 0.86 | 0.87 | 57 | 7 $\beta$-strands |
| | | 2 | 0.84 | 0.86 | 90 | 7 $\beta$-strands |
| PD-L1 | 5o45 | 1 | 0.91 | 0.91 | 45 | Double helix |
| | | 2 | 0.90 | 0.90 | 97 | 4 helices |
| | | 3 | 0.90 | 0.90 | 61 | 3 helices |
| MDM2 | 1ycr | 1 | 0.78 | 0.88 | 14 | Single helix |
| GLP-1R | 3iol | 1 | 0.86 | 0.91 | 20 | Single helix |
| KRasG12C | 8x6r | 1 | 0.85 | 0.69 | 78 | 4 helices |
| IL7$-$R$\alpha$ | 3di3 | 1 | 0.78 | 0.88 | 28 | Single helix |

**Table 3**: Protein target binder design analysis. The table presents various protein targets with their reference PDB structures and designed binders. For each design, AlphaFold3 (AF3) complex prediction scores (iPTM and pTM), amino acid length, and secondary structure characteristics of the designed binders are shown.

Designing high-affinity binders is one of the most critical and practically valuable challenges in *de novo* protein design. Traditionally, binder design has relied on borrowing natural complexes and iteratively introducing sequence mutations in hotspot regions, guided by physical principles, to achieve improved binders. These methods are significantly influenced by natural sequences, limiting the scope of mutations and innovation. They often require thousands of experiments to screen for a few useful binders, resulting in very low success rates.

| Type | Reference | iPTM | pTM | Length | Secondary Structure | MD | Name |
|---|---|---|---|---|---|---|---|
| VEGF | 1bj1 | 0.83 | 0.84 | 80 | 3 helices + 3 $\beta$-sheets | -15 | VE1 |
| VEGF | 1bj1 | 0.79 | 0.82 | 110 | 8 $\beta$-strands | -48 | VE2 |
| VEGF | 1bj1 | 0.83 | 0.84 | 114 | 9 $\beta$-strands | -35 | VE3 |
| VEGF | 1bj1 | 0.85 | 0.86 | 58 | 3 helices | -45 | VE4 |
| VEGF | 1bj1 | 0.85 | 0.86 | 95 | 6 $\beta$-sheets | -36 | VE5 |
| VEGF | 1bj1 | 0.81 | 0.83 | 51 | 3 helices | -63 | VE6 |
| VEGF | 1bj1 | 0.82 | 0.83 | 116 | 8 $\beta$-strands | -50 | VE7 |
| VEGF | 1bj1 | 0.84 | 0.85 | 95 | 6 $\beta$-sheets | -45 | VE8 |
| VEGF | 1bj1 | 0.82 | 0.84 | 58 | 3 helices | -62 | VE9 |

**Table 4**: VEGF (Vascular Endothelial Growth Factor) binder design analysis. The table presents 12 designed binders with their AlphaFold3 complex prediction scores (iPTM and pTM), amino acid lengths, secondary structure characteristics, and molecular dynamics (MD) scores. All designs target VEGF using the 1bj1 PDB structure as reference.

Affinity Verification - Summary

**VEGF**

| Name | KD(M) |
|---|---|
| VE1 | 3.5E-5 |
| VE3 | 1.7E-4 |
| VE4 | 4.96E-5 |
| VE5 | 6.99E-5 |
| VE6 | 1.28E-4 |
| VE7 | 1.01E-4 |
| VE8 | 6.17E-5 |
| VE9 | 3.8E-4 |

**PDL1**

| Name | KD(M) |
|---|---|
| PDL1_1 | 1.962E-4 |
| PDL1_2 | 3.974E-5 |
| PDL1_3 | 9.508E-5 |

**COVID SRC2RBD**

| Name | KD(M) |
|---|---|
| Covid_1 | 5.944E-5 |
| Covid_2 | 7.20E-5 |
| Covid_3 | 7.371E-5 |
| Covid_4 | 1.989E-6 |
| Covid_5 | 5.401E-5 |

**Fig. 7**: Wet experimental affinity results for three targets: PDL1, COVIDSRC2RBD, and VEGF

Recent approaches, such as RFdiffusion and other deep learning-based methods, have used generative models to produce binders with some success. However, models like RFdiffusion still require the specification of key hotspots. A rational model that has learned the interactions between protein complexes should be capable of automatically identifying critical regions, as these areas typically exhibit significant structural differences (e.g., forming strong non-covalent interactions), which are essential for creating effective binders. Therefore, we believe that binder design should be more end-to-end.

Based on the aforementioned foundational models, we fine-tuned our approach using complexes and PDBBind data. We used the structural and sequence features of the target chain as input to directly generate the diffused chain. We tested our model on various typical binder structures and some high-value but highly challenging binders.

For any given target, we input the model-generated diffused chain into Protein MPNN for sequence filling. Then, we input the native target sequence and the diffused chain's sequence into AF2 to predict the complex structure. We classified a design as successful if it had an AF2 pAE of interaction between the binder and target of less than 10 (a strong indicator of design success), an RMSD between the designed binder and the AF2 prediction of less than 1Å, and an AF2 plDDT greater than 80.

#### J.1 Design for PD-L1

Programmed Death-Ligand 1 (PD-L1) is a key immune checkpoint protein that suppresses T-cell activity, enabling cancer cells to evade immune surveillance. Using the PD-L1 structure from PDB `5o45` as the target, we designed 20 initial candidates. Based on predicted alignment error (pAE), three structurally diverse designs (5o45_1, 5o45_4, 5o45_18) were selected for AlphaFold 3 (AF3) evaluation.

The best-performing binder, 5o45_18, achieved an initial ipTM/pTM of 0.75/0.79. After a round of secondary sequence optimization, the final design reached ipTM = 0.88 and pTM = 0.87. This binder was validated in *in vitro* experiments, demonstrating high expression, solubility, and binding affinity.

#### J.2 Design for MDM2

MDM2 is a key negative regulator of the tumor suppressor p53, promoting its degradation through direct binding. Targeting MDM2 can help restore p53 activity in cancers. Using the MDM2 structure from PDB `1ycr`, we designed 20 binder candidates.

Seven cases with pLDDT $>$ 70 were selected for AF3 evaluation. The best binder, 1ycr_3, achieved ipTM = 0.78 and pTM = 0.87. No further optimization was performed.

#### J.3 Design for SARS-CoV-2 Spike RBD

The receptor-binding domain (RBD) of the SARS-CoV-2 spike protein mediates viral entry via ACE2 binding. Using the RBD from PDB `7RBY` as the target, we generated 50 initial binders. Sequence completion and ESMFold folding were performed, followed by selection of the top 20 by PLDDT.

From these, five structurally diverse designs were refined through two iterations. All five achieved ipTM and pTM values approaching 0.8. Despite some cases having $ipTM > 0.7$ only, we proceeded directly to molecular dynamics simulations and *in vitro* experiments.

#### J.4 Design for GLP-1R

GLP-1R is a class B GPCR involved in glucose regulation. Binding of GLP-1 to its receptor stimulates insulin secretion and is a common therapeutic strategy. Using the GLP-1R complex structure from PDB `3iol` as reference, we generated 20 initial designs.

Without secondary optimization, the top candidate 3iol_18 achieved ipTM = 0.86 and pTM = 0.91 in AF3 evaluation.

#### J.5 Design for HIV GP120

gp120 is the HIV envelope protein that binds to CD4 on host cells, initiating viral entry. We used two PDB structures—`5cd5` and `2ny3`—as references. For 5cd5, 100 designs were generated and filtered by $PLDDT > 70$ and $pAE < 5$. Three were selected for AF3 refolding. After secondary sequence optimization on 5cd5_96, we obtained a binder with ipTM = 0.88 and pTM = 0.87.

Using 2ny3, we selected 8 diverse designs for AF3 prediction. Two underwent refinement and achieved ipTM/pTM values above 0.8: 2ny3_63 (0.84/0.86) and 2ny3_75 (0.86/0.87).

#### J.6 Design for KRasG12

KRasG12C is a common oncogenic mutation found in several cancers and is considered a difficult target. Using the mutant KRas structure from PDB `8x6r`, we generated 100 initial binders. Based on $pLDDT > 70$, $pAE < 3$, and structural diversity, three cases (8x6r_40, 8x6r_43, 8x6r_74) were selected.

After sequence refinement, 8x6r_74 achieved ipTM = 0.85 and pTM = 0.69. The other two designs had lower structural compatibility.

Additionally, we designed four binders targeting the G12D epitope based on the 7rpz protein, as shown in SI Table 3.

#### J.7 Design for IL-7R$\alpha$

IL-7R$\alpha$ activates T cell development, proliferation, and survival signals by binding to IL-7. It plays a critical role in early T cell development and the maintenance of peripheral T cells. Aberrant expression or dysfunction of IL-7R$\alpha$ is associated with various immune-related diseases, such as immunodeficiency, lymphoma, and autoimmune disorders. Based on the target region in the 3di3 protein, we designed one binder, resulting in a design outcome.

#### J.8 Design for VEGF

Vascular Endothelial Growth Factor (VEGF) is a central regulator of angiogenesis and a key therapeutic target in cancer and ocular diseases. Using the PDB structure 1BJ1 as reference, we generated 100 initial designs with lengths between 50 and 200 residues.

The top 20 diverse structures (by PLDDT) were refined for up to two iterations. We ultimately obtained 9 high-confidence *in silico* binders. These were validated through *in vitro* experiments, confirming strong expression and affinity.

#### J.9 Gromacs MD rmsd for Designed binders

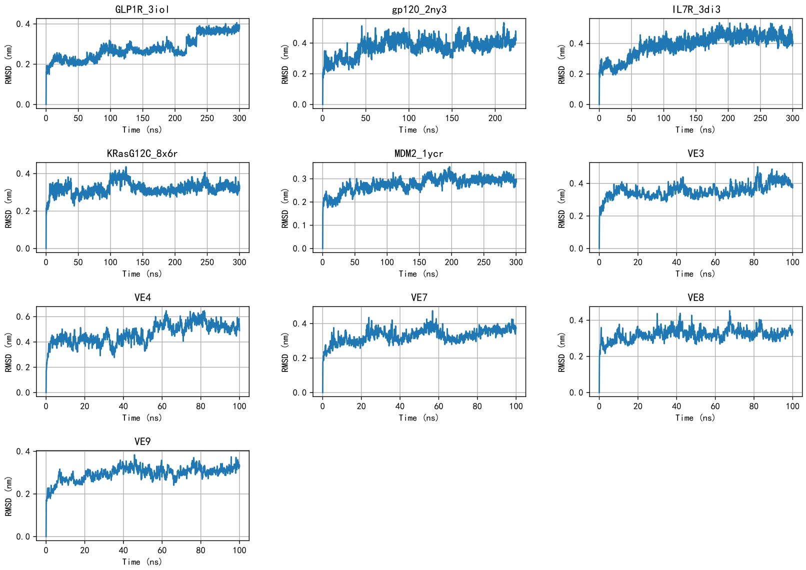


**Fig. 8**: Root Mean Square Deviation (RMSD) of protein structures over time under different conditions. Each subplot represents the RMSD trajectory for a specific simulation condition, with time in nanoseconds (ns) and RMSD in nanometers (nm). RMSD values indicate the structural deviation from the reference structure, where higher values correspond to greater structural changes.

In molecular dynamics simulations, we evaluated the structural stability of the protein by calculating the Root Mean Square Deviation (RMSD). As shown in Figure 8, we compared the RMSD variations under different conditions. The RMSD was calculated based on the average deviation of protein backbone atoms from their initial positions, with higher values indicating more significant structural changes. The

RMSD trajectories reveal distinct patterns of structural evolution under various conditions, providing insights into the conformational dynamics and structural stability of the protein. This analysis allows us to understand how different environmental conditions affect the protein's structural integrity and conformational changes.
