## Supplementary material for "Robust and Reliable *de novo* Protein Design: A Flow-Matching-Based Protein Generative Model Achieves Remarkably High Success Rates": SICD

Circular Dichroism (CD) Spectroscopy of PDL1 binder

PDL1-1

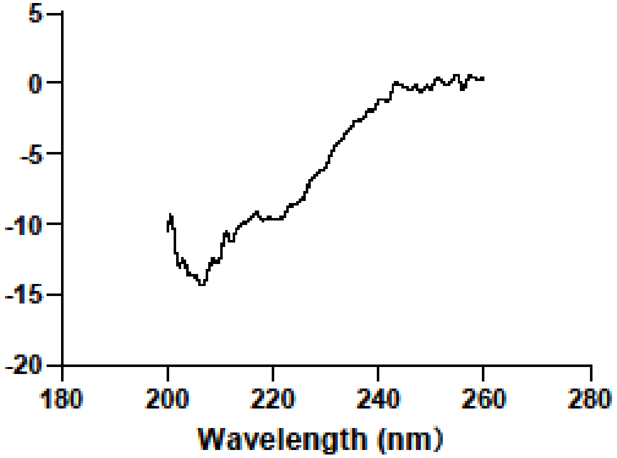


PDL1-2

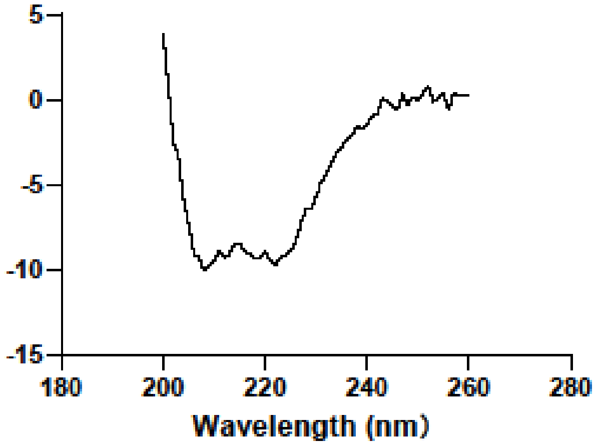


PDL1-3

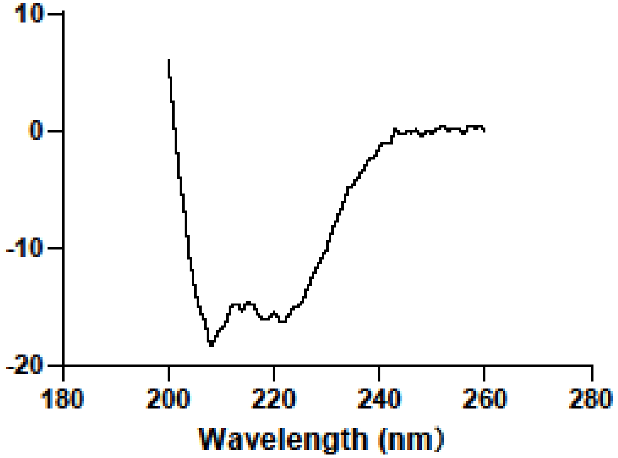

### Circular Dichroism (CD) Spectroscopy of Covid SRC2RBD binder

#### Covid-1

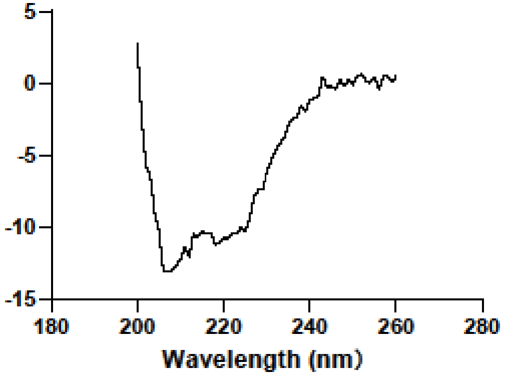


#### Covid-2

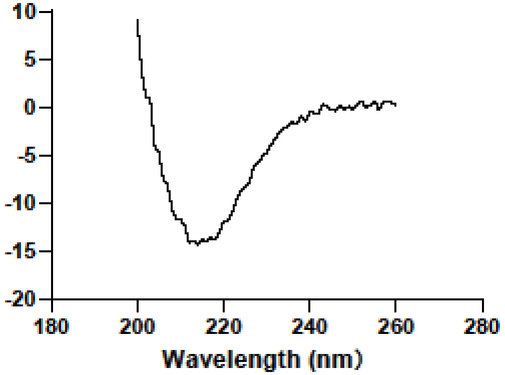


#### Covid-3

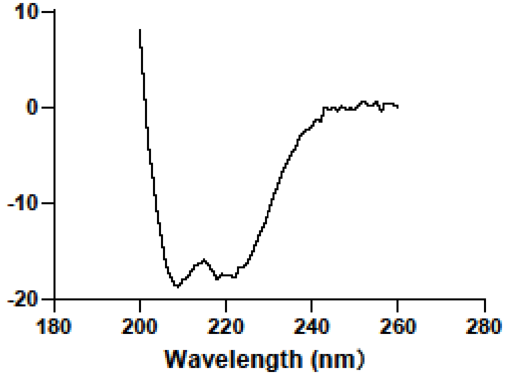


#### Covid-4

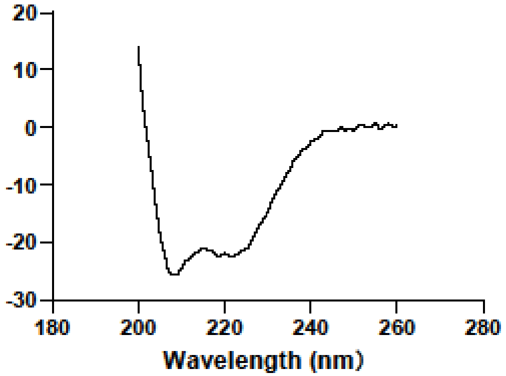


#### Covid-5

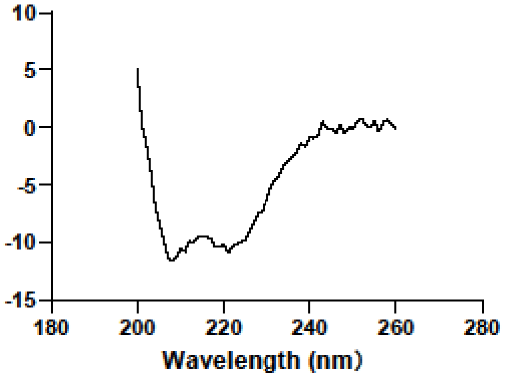

### Circular Dichroism (CD) Spectroscopy of VEGF binder

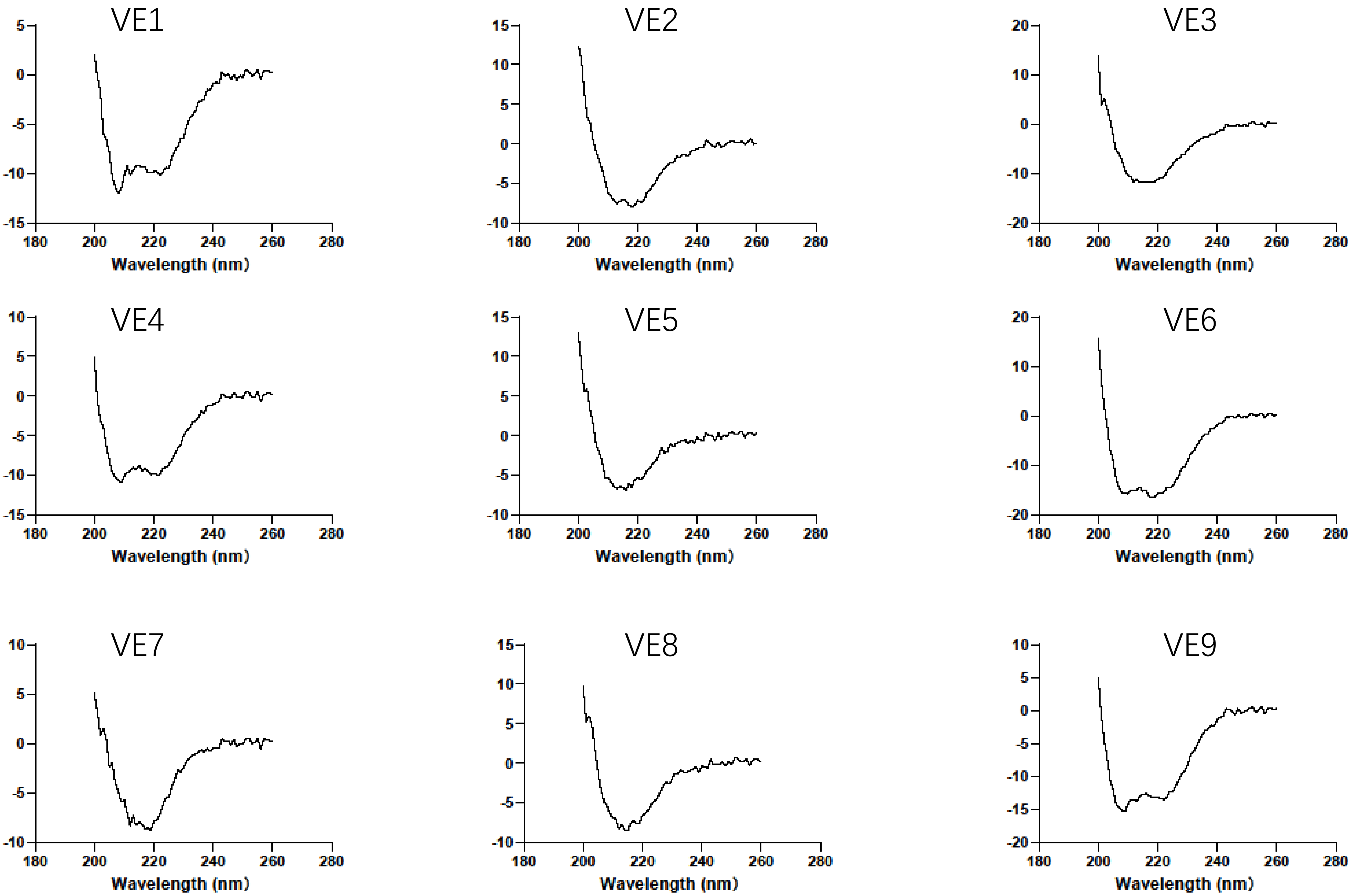
