## Supplementary figures and images for "Robust and Reliable *de novo* Protein Design: A Flow-Matching-Based Protein Generative Model Achieves Remarkably High Success Rates"

### SISDS

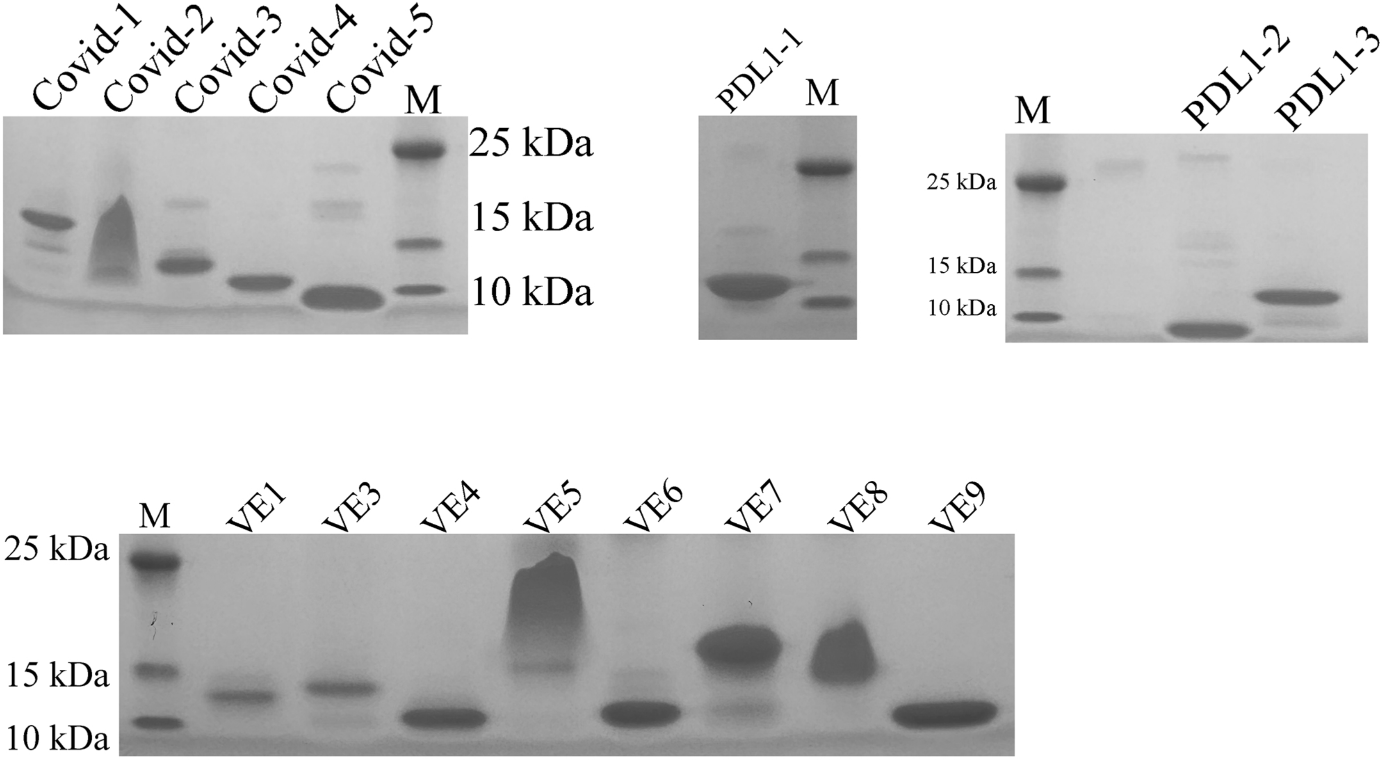

### SITM

#### Thermal Melting (TM) Plot of Binders Designed by OriginFlow

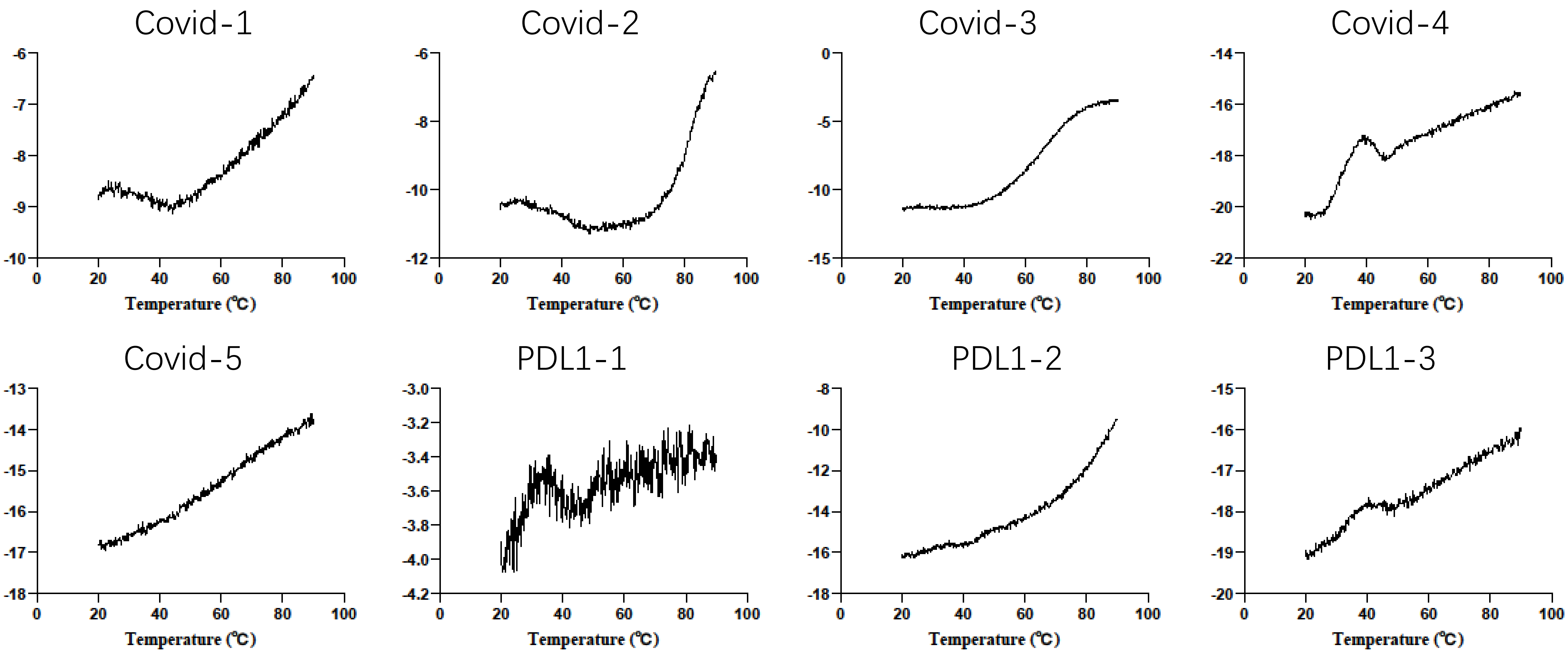

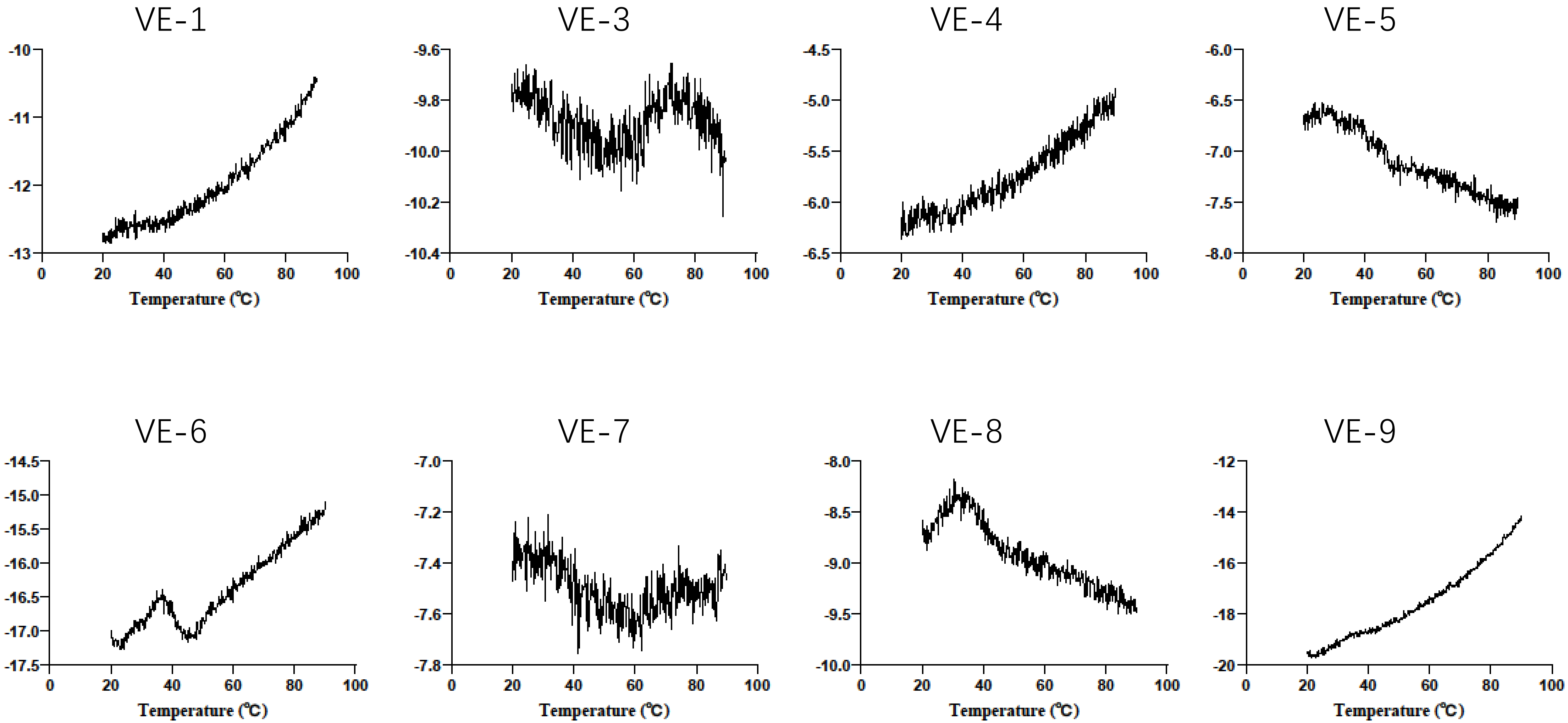
