## Supplementary material for "Robust and Reliable *de novo* Protein Design: A Flow-Matching-Based Protein Generative Model Achieves Remarkably High Success Rates": SISPR

### SPR Results of Target Designs by OriginFlow

Affinity Validation - Summary

**VEGF**

| Name | KD(M) |
|---|---|
| VE1 | 3.5E-5 |
| VE3 | 1.7E-4 |
| VE4 | 4.96E-5 |
| VE5 | 6.99E-5 |
| VE6 | 1.28E-4 |
| VE7 | 1.01E-4 |
| VE8 | 6.17E-5 |
| VE9 | 3.8E-4 |

**RBD**

| Name | KD(M) |
|---|---|
| Covid-1 | 5.94E-05 |
| Covid-2 | 7.20E-05 |
| Covid-3 | 7.37E-05 |
| Covid-4 | 1.99E-06 |
| Covid-5 | 5.40E-05 |

**PDL1**

| Name | KD(M) |
|---|---|
| PDL1-1 | 1.962E-4 |
| PDL1-2 | 9.508E-5 |
| PDL1-3 | 3.974E-5 |

### SPR Experimental Results of RBD Ligands

#### Covid-1

Report table

| KD (M) | Rmax (RU) | offset (RU) | Chi² (RU²) |
|---|---|---|---|
| 5.944E-5 | | | 5.55 |
| | 106.8 | 8.292 | |

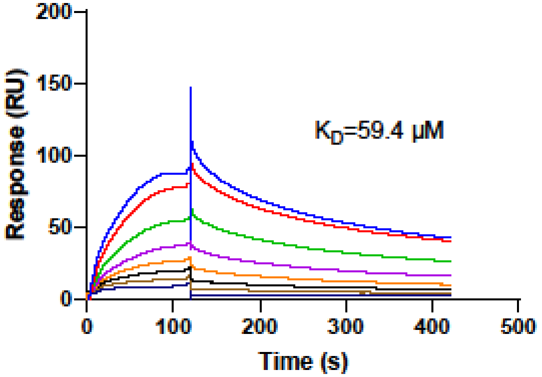


#### Covid-2

Report table

| KD (M) | Rmax (RU) | offset (RU) | Chi² (RU²) |
|---|---|---|---|
| 7.200E-5 | | | 29.7 |
| | 230.6 | 14.77 | |

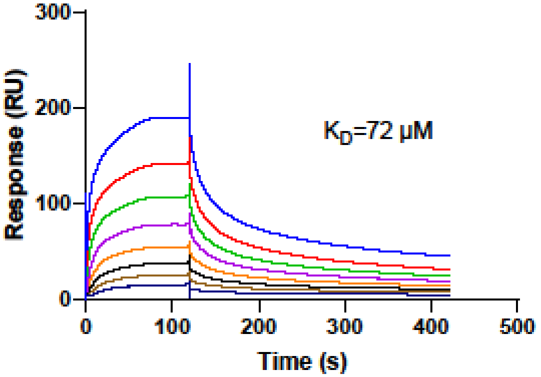


#### Covid-3

Report table

| KD (M) | Rmax (RU) | offset (RU) | Chi² (RU²) |
|---|---|---|---|
| 7.371E-5 | | | 18.9 |
| | 158.4 | 8.609 | |

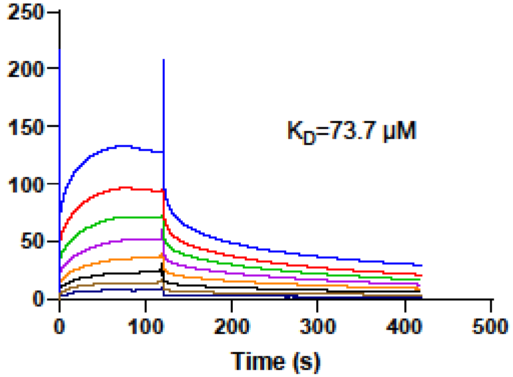


#### Covid-4

Report table

| KD (M) | Rmax (RU) | offset (RU) | Chi² (RU²) |
|---|---|---|---|
| 1.989E-6 | | | 0.271 |
| | 17.05 | -2.218 | |

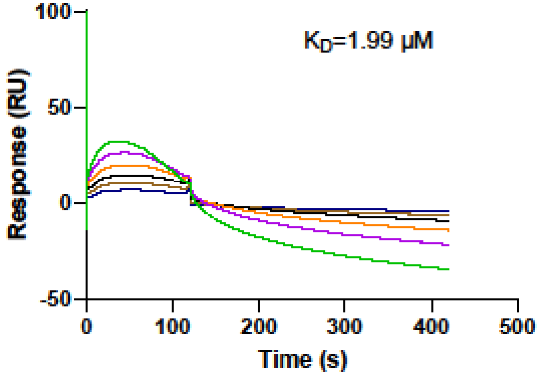


#### Covid-5

Report table

| KD (M) | Rmax (RU) | offset (RU) | Chi² (RU²) |
|---|---|---|---|
| 5.401E-5 | | | 4.02 |
| | 61.65 | 1.768 | |

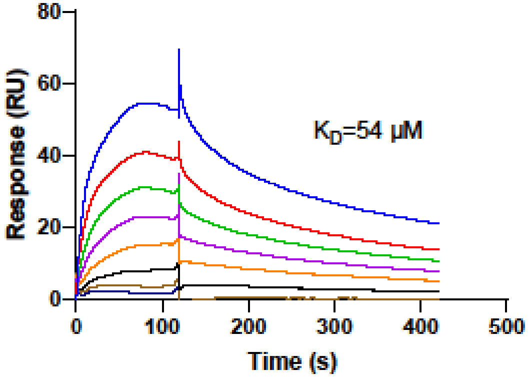


| Name | KD(M) |
|---|---|
| Covid-1 | 5.94E-05 |
| Covid-2 | 7.20E-05 |
| Covid-3 | 7.37E-05 |
| Covid-4 | 1.99E-06 |
| Covid-5 | 5.40E-05 |

### SPR Experimental Results of PDL1 Ligands

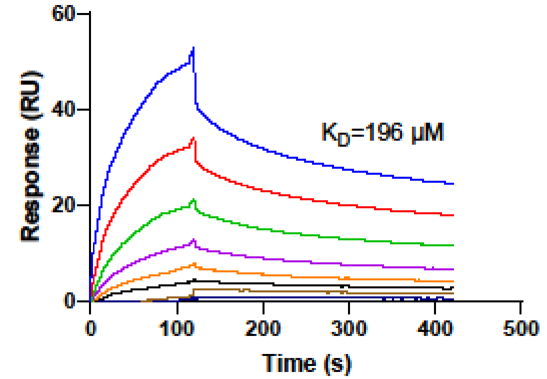


Report table

| KD (M) | Rmax (RU) | offset (RU) | Chi² (RU²) |
|---|---|---|---|
| 1.962E-4 | | | 0.869 |
| | 99.74 | 0.1213 | |

PDL1-1

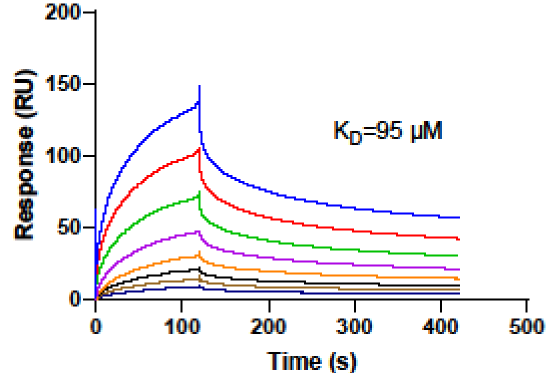


Report table

| KD (M) | Rmax (RU) | offset (RU) | Chi² (RU²) |
|---|---|---|---|
| 9.508E-5 | | | 2.72 |
| | 187.5 | 6.822 | |

PDL1-2

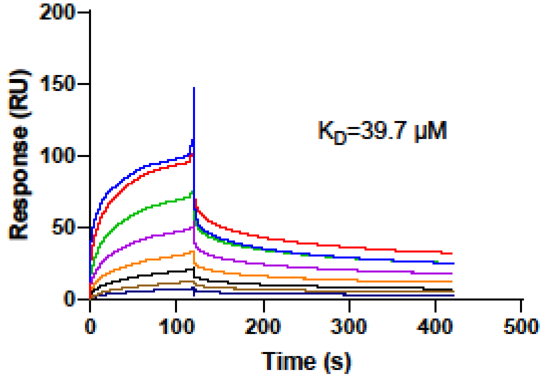


Report table

| KD (M) | Rmax (RU) | offset (RU) | Chi² (RU²) |
|---|---|---|---|
| 3.974E-5 | | | 7.46 |
| | 126.7 | 2.158 | |

PDL1-3

| Name | KD(M) |
|---|---|
| PDL1-1 | 1.962E-4 |
| PDL1-2 | 9.508E-5 |
| PDL1-3 | 3.974E-5 |

### Affinity Validation of VEGF and Its Binders

## VE1

Parameters table

| KD (M) | SE(KD) | Rmax (RU) | SE(Rmax) | offset (RU) | SE(offset) |
|---|---|---|---|---|---|
| 3.501E-5 | 2.1E-6 | | | | |
| | | 1171.8 | 23 | 9.8 | 6.8 |

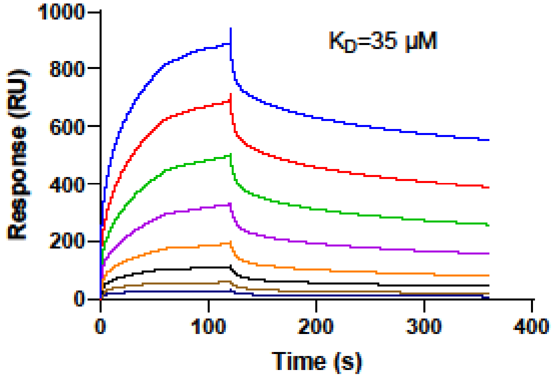


VE2 was excluded from testing due to the formation of fibrous precipitates in the protein.

## VE3

Parameters table

| KD (M) | SE(KD) | Rmax (RU) | SE(Rmax) | offset (RU) | SE(offset) |
|---|---|---|---|---|---|
| 1.701E-4 | 3.2E-5 | | | | |
| | | 632.0 | 60 | 4.0 | 5.2 |

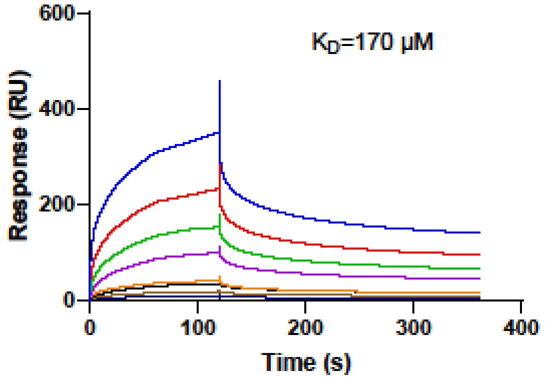


## VE4

Parameters table

| KD (M) | SE(KD) | Rmax (RU) | SE(Rmax) | offset (RU) | SE(offset) |
|---|---|---|---|---|---|
| 4.966E-5 | 6.7E-6 | | | | |
| | | 1792.5 | 67 | 69.3 | 30 |

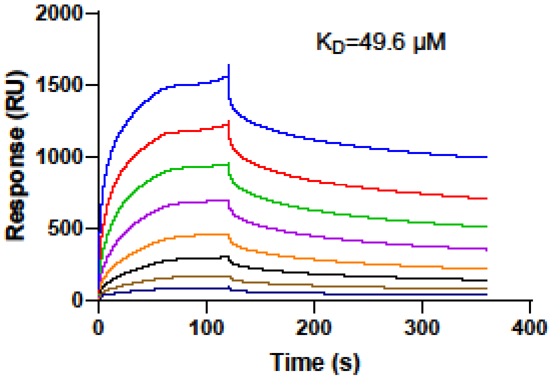


## VE5

Parameters table

| KD (M) | SE(KD) | Rmax (RU) | SE(Rmax) | offset (RU) | SE(offset) |
|---|---|---|---|---|---|
| 6.996E-5 | 5.0E-6 | | | | |
| | | 630.8 | 15 | 15.6 | 4.3 |

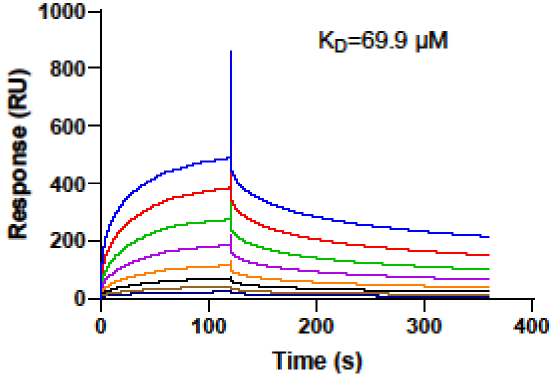


## VE6

Parameters table

| KD (M) | SE(KD) | Rmax (RU) | SE(Rmax) | offset (RU) | SE(offset) |
|---|---|---|---|---|---|
| 1.279E-4 | 2.1E-5 | | | | |
| | | 1378.0 | 100 | 40.9 | 13 |

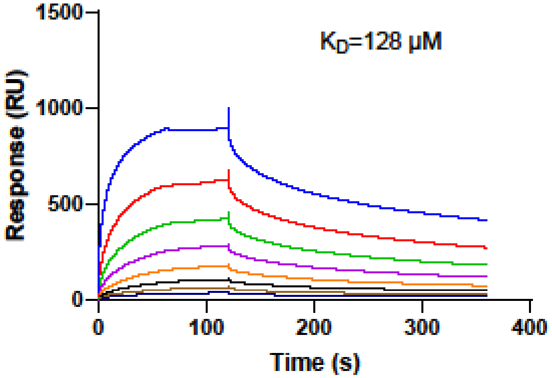

## VE7

Parameters table

| KD (M) | SE(KD) | Rmax (RU) | SE(Rmax) | offset (RU) | SE(offset) |
|---|---|---|---|---|---|
| 1.014E-4 | 1.3E-5 | | | | |
| | | 601.9 | 31 | 17.0 | 5.6 |

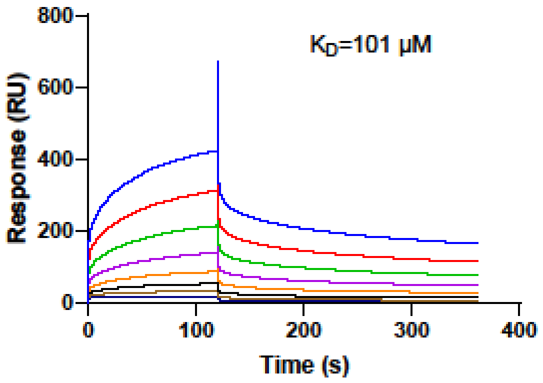


## VE8

Parameters table

| KD (M) | SE(KD) | Rmax (RU) | SE(Rmax) | offset (RU) | SE(offset) |
|---|---|---|---|---|---|
| 6.173E-5 | 6.6E-6 | | | | |
| | | 593.2 | 19 | 24.2 | 6.7 |

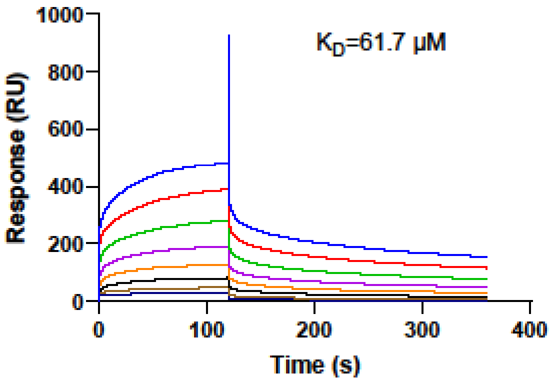


## VE9

Parameters table

| KD (M) | SE(KD) | Rmax (RU) | SE(Rmax) | offset (RU) | SE(offset) |
|---|---|---|---|---|---|
| 3.804E-4 | 1.4E-4 | | | | |
| | | 337.7 | 86 | -3.0 | 2.1 |

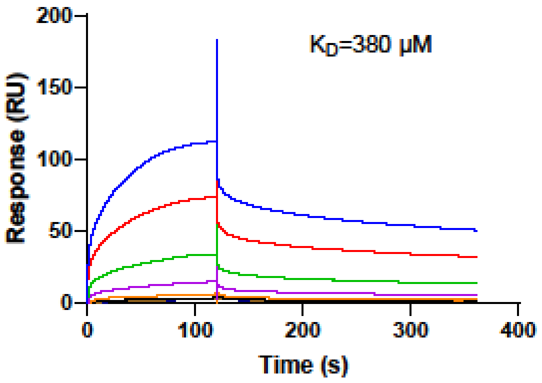
